# Amyloid precursor protein localises to ependymal cilia in vertebrates and is required for ciliogenesis and brain development in zebrafish

**DOI:** 10.1101/2021.06.02.446706

**Authors:** Jasmine Chebli, Maryam Rahmati, Tammaryn Lashley, Birgitta Edeman, Anders Oldfors, Henrik Zetterberg, Alexandra Abramsson

## Abstract

Amyloid precursor protein (APP) is ubiquitously expressed in human, mice and in zebrafish. In zebrafish, there are two orthologues, Appa and Appb. Interestingly, some cellular processes associated with APP overlap with cilia-mediated functions. Whereas the localization of APP to primary cilia of *in vitro*-cultured cells has been reported, we addressed the presence of APP in motile and in non-motile sensory cilia and its potential implication for ciliogenesis using zebrafish, mouse, and human samples. We report that Appa and Appb are expressed by ciliated cells and become localized at the membrane of cilia in the olfactory epithelium, otic vesicle and in the brain ventricles of zebrafish embryos. App in ependymal cilia persisted in adult zebrafish and was also detected in mouse and human brain. Finally, we found morphologically abnormal ependymal cilia and smaller brain ventricles in *appa^-/-^appb^-/-^* mutant zebrafish. Our findings demonstrate an evolutionary conserved localisation of APP to cilia and suggest a role of App in ciliogenesis and cilia-related functions.

## Introduction

Amyloid precursor protein (APP) is a ubiquitously expressed type-1 transmembrane protein that, together with the APP-like protein 1 and -2 (APLP1, APLP2), comprises the *APP* gene family. In addition to their various splice forms, they are all post-translationally modified through proteolytic processing (1). Although the physiological relevance of the fragments generated is not fully understood, one of these, the amyloid-beta peptide (Aβ) originating from the transmembrane domain of the APP protein, is the main component of brain amyloid plaques in Alzheimer’s disease (AD) (1, 2). Beyond its pathological involvement, studies on APP have revealed essential physiological functions including neurogenesis (3, 4), neurite outgrowth (5, 6), adhesion properties (6, 7), synapse formation (8), and neuronal migration (6, 9, 10). Nevertheless, the involvement of each APP family member in these processes remains unclear, since redundancy makes it difficult to unravel the contribution of a specific protein (11). Although the molecular mechanisms behind the APP-related processes are yet to be determined, accumulating evidence support that APP orchestrates cellular processes through receptor-like interactions with both inter- and intra-cellular signaling molecules (6).

The cilium is a highly conserved organelle across species, which contributes to a wide range of cellular processes (12). Cilia can broadly be categorized into motile and non-motile. Non-motile cilia include primary cilia, which are ubiquitously expressed on most cells as a single short antenna-like structure, and sensory cilia, that are only expressed by specific cells. Primary cilia are enriched in receptors and sites for inter-cell signaling transduction and are notably implicated in cell division, autophagy, midbrain development, memory and learning abilities (13). As for the sensory cilia, they are notably found in the otic vesicle as stereocilia and kinocilia. Motile cilia are present on cells involved in fluid movement including the epithelium of the respiratory tract and the ependymal layer of the brain ventricles. Ependymal cells are derived from radial glial cells and when fully differentiated are decorated with tufts of motile cilia anchored with roots at the apical cellular membrane (12, 14). The coordinated periodic beating of the cilia participate in the generation of cerebrospinal fluid (CSF) flow within ventricle cavities (15). Circulation of CSF is believed to facilitate transfer of signaling molecules and removal of metabolic waste products to prevent accumulation of neurotoxic residues in the brain parenchyma (16–18).

There are several findings supporting a connection between APP and cilia. First, part of the wide range of cilia-mediated functions overlap with processes linked to APP, *e.g.*, cognitive impairment (19), differentiation of neurons (20), formation of corpus callosum (19, 21), neuronal migration (22–24) and sensing of guidance molecules (25). Second, overexpression of APP impairs primary cilia both in a mouse AD model and in individuals with Down syndrome, harboring three copies of *APP* (26, 27). The latter is also associated with decreased CSF flow and accumulation of CSF (hydrocephalus), two phenotypes commonly associated with defects in motile cilia (28). Finally, APP has been shown to localize to primary cilia *in vitro* and Aβ exposure results in reduced cilia length (29). Taken together, these findings warrant further investigations of the role of APP in both motile and non-motile cilia.

In the present study, we address the presence of APP in motile and non-motile (sensory) cilia and its possible functions using zebrafish, mouse and human samples. We found that the zebrafish App homologues are expressed by ciliated cells and become localized at the membrane of cilia in the otic vesicle, the nasal epithelium, and the brain ventricles of zebrafish embryos. The presence of App in ependymal cilia persisted in adult zebrafish and was also detected in the ependymal cells of mouse and human brains. In addition, we show that zebrafish embryos with mutations in both *app* paralogues (*appa^-/-^appb^-/-^)* have morphologically abnormal ependymal cilia and smaller brain ventricles compared with wild-type siblings.

## Results

### Appa and appb mRNA expression patterns at the brain ventricular limits

The zebrafish *app* genes, *appa* and *appb,* are expressed in the CNS, and have both distinct and shared expression patterns (7, 30). Due to the lack of specific antibodies, we used fluorescent whole mount *in situ* hybridization to increase the cellular resolution of *appa* and *appb* mRNA expression in areas with motile cilia on 30 hpf wild-type larvae zebrafish (***Figure 1***). Consistent with previous studies, we observed *appa* mRNA expression in the lens, the olfactory bulb and epithelium, in the trigeminal ganglia and in the otic vesicle. (***Figure 1C***). Similarly, the *appb* mRNA expression signal corroborated previous data on *appb* mRNA expression (30) in the olfactory and otic vesicle epithelia (***Figure 1H***).

**Figure 1.**
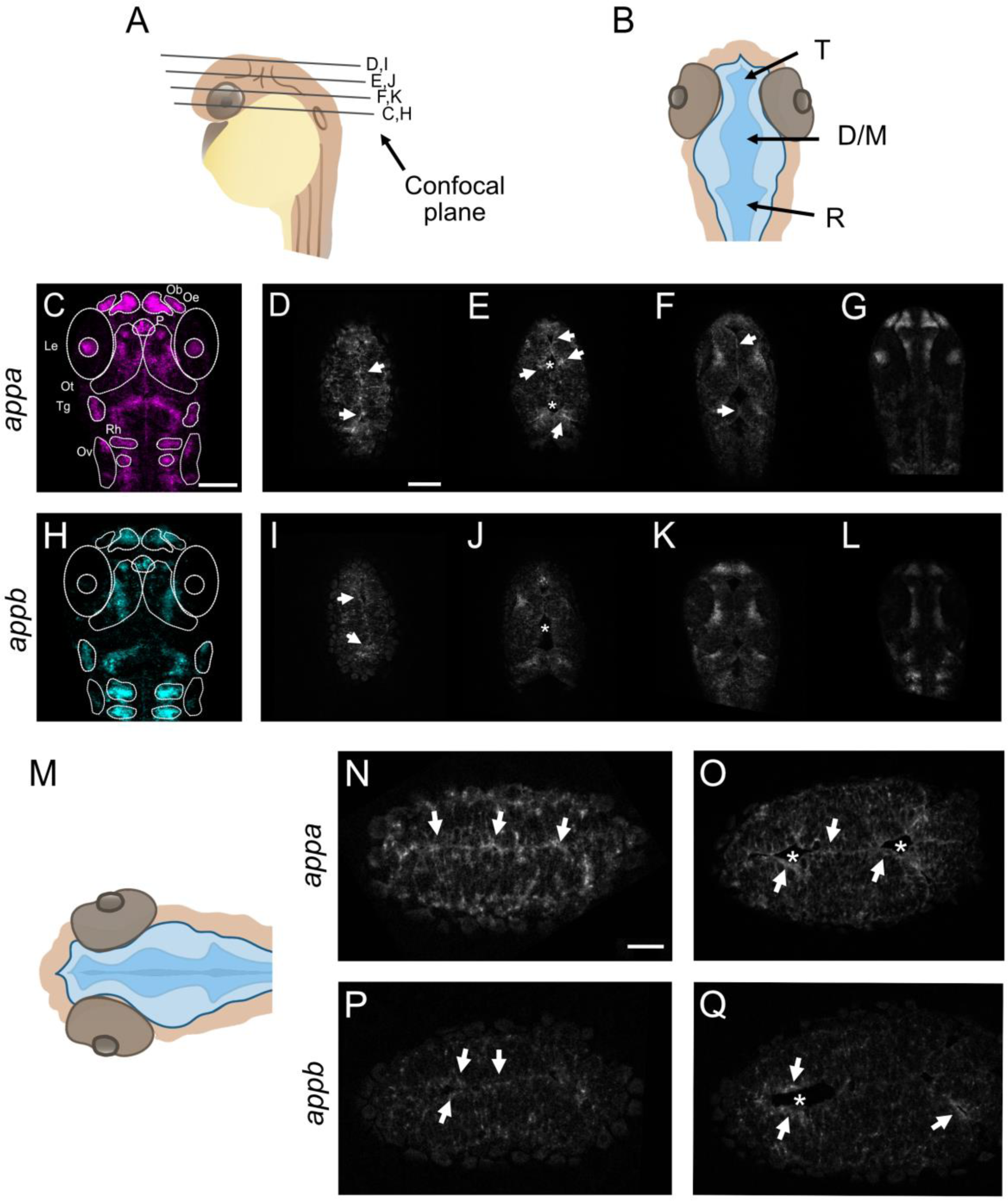
Expression of appa and appb mRNA in zebrafish larvae. Expression pattern of *appa* and *appb* mRNA. (**A,B**) Schematic representations of head and ventricle morphology of a 30 hpf zebrafish larvae, lateral (**A**) and dorsal (**B**) view. (**C,H**) Whole-mount fluorescent *in situ* of *appa* (**C**) and *appb* (**H**) in 30 hpf WT zebrafish larvae. Single focal planes, dorsal to ventral, of whole-mount larvae of *appa* (**D-G**) and *appb* (**I-L**) probe. (**M**) Schematic view of focal plane of the dorsal area of the brain ventricle. (**N-Q**) Single focal plane at high magnification (40x) of *appa* (**N,O**) and *appb* (**P,Q**) probes. T= telencephalic ventricle, D/M= diencephalic/mesencephalic ventricle, R= rhombencephalic ventricle, Ob= olfactory bulb, Oe= olfactory epithelium, P= pituitary gland, Le= lens, Ot= optic tectum, Tg= trigeminal ganglia, Rh= rhombomeres, Ov= otic vesicle. Magnification: (**C-L**)= 20x, (**N-Q**)= 40x. Scale bar: (**C**)=100µm, (**D**)= 50µm, (**N**)= 25µm. * indicates ventricular space and arrows highlight expression.

In addition, both *appa (****Figures 1C-G and high magnification Figures 1N,O)*** and *appb* (***Figures 1H-L, P,Q***) mRNA signals labelled cells lining the diencephalic ventricle both in the dorsal and ventral areas. Negative controls did not show any specific signal (***Supplementary file 1***). Together, these results show expression of *appa* and *appb* in areas with ciliated cells, including cells lining brain ventricles, otic vesicle and olfactory organ, thus suggesting a possible role of App in cilia formation and function.

### App protein is localized to cilia of the olfactory sensory neurons and otic vesicle in zebrafish larvae

The expression of both *appa* and *appb* in ciliated cells made us ask if the proteins become distributed out to the cilia. The zebrafish olfactory epithelium and the otic vesicle comprise ciliated cells and are regions where both *appa* and *appb* mRNAs are expressed. To address if Appa and Appb become localized to these cilia, we performed immunofluorescent staining on zebrafish larvae.

### Olfactory sensory neuron cilia

We used the Y188 antibody, binding to a conserved epitope in the C-terminal end of human, mouse, and zebrafish App (***Figure 6C***), in combination with the anti-acetylated tubulin antibody, labelling microtubule structures of cilia. Immunofluorescent co-labelling detected a punctate App signal in the heavily ciliated olfactory epithelium area at 30 hpf (***Figure 2A***). However, while the resolution of the images did not allow distinction between each cilium, App signal seemed to localize to most of them. In addition to the cilium, App expression was also found at the base of these motile cilia (***Figure 2A’***).

**Figure 2.**
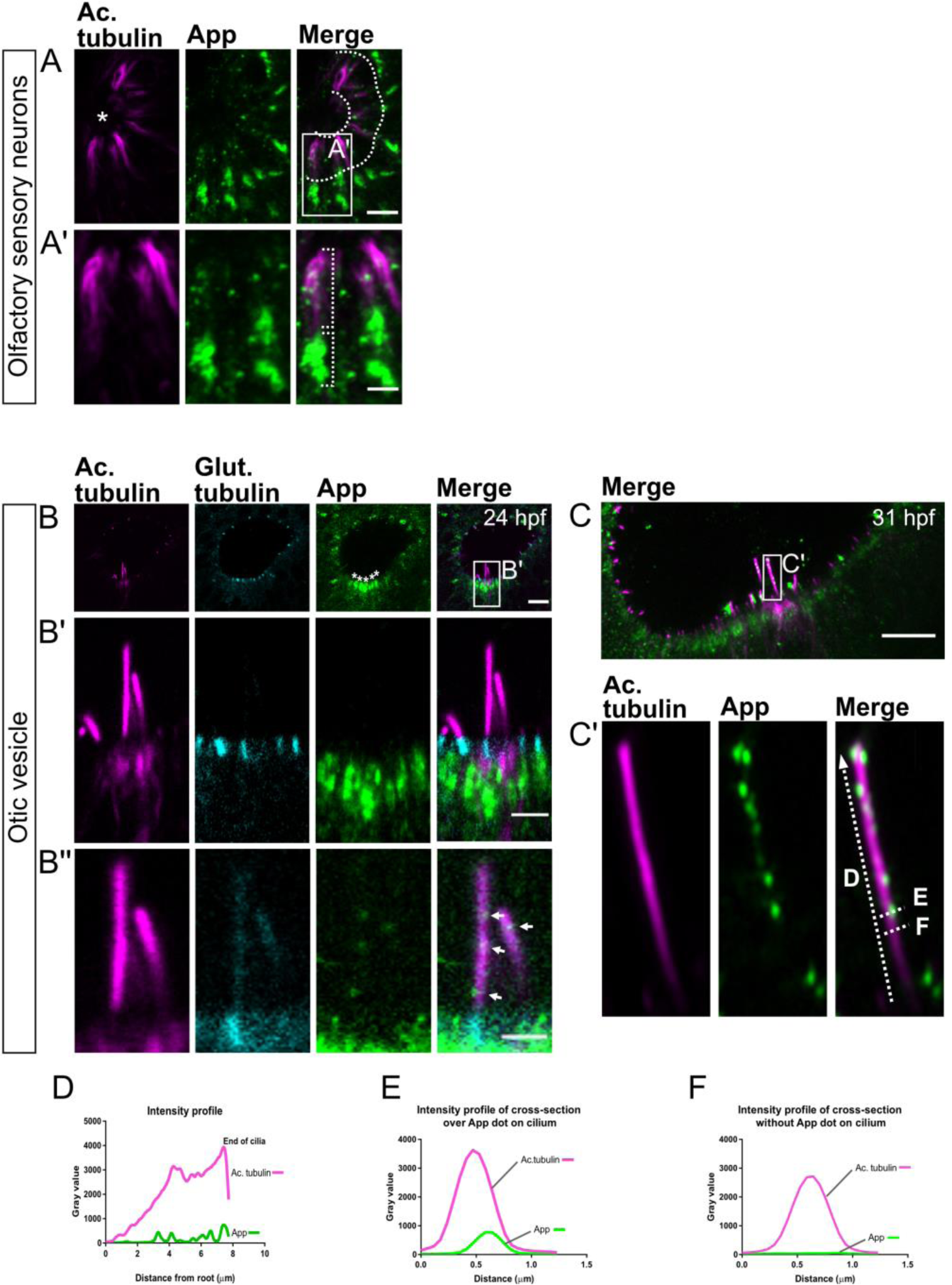
App protein is localized to non-primary cilia in zebarfish larvae. Localization of App protein to cilia of the olfactory sensory neurons and otic vesicle in 31 hpf larvae. Cilia as shown by immunostaining for acetylated tubulin (magenta) and App (green) of the olfactory sensory neurons in the nose epithelium (**A**) and the otic vesicle (**B-C**). In (**A**), dotted lines demarcate the cilia from the nasal cavity (see asterisk). (**A’**) App (green) is found along the cilia and accumulating at their base. Otic vesicle of 24 hpf (**B**) and 31 hpf larvae (**C**). In (**B**), glutamylated tubulin (cyan) highlights the base of the cilia outlined by acetylated tubulin staining (magenta). (**B**) Overview of the kinocilia and stereocilia of the otic vesicle. The white asterisks indicate accumulation of App (green) at the base of the cilia bundles. (**B’**) Magnification of cilia outlined in (**B**). (**B’’**) Increased intensity of the green channel to detect App (arrows) in kinocilia. Otic vesicle in 31hpf zebrafish larvae (**C**) with close-up (**C’**) showing App puncta (green) along the kinocilia. Intensity profiles of acetylated tubulin (magenta) and App (green) staining from the kinocilia (**D-F**). In (**D**), the intensity profile of the whole length of the kinocilium is plotted whereas profiles of cross-section lines are plotted with a visible App puncta (**E**) and without (**F**). The dotted lines (**C’**) indicate the kinocilium and cross-sections. Magnification: (**A-C**)= 40x. Scale bar: (**A**)= 5 µm, (**B**)= 10 µm, (**B’**)= 4 µm, (**B’’**)= 2 µm, (**C**)= 10µm.

### Otic vesicle cilia

Similar to the olfactory neurons, high accumulation of App was noted at the base of the cilia in the otic vesicle. In zebrafish, hair cells of the otic vesicle have two types of cilia, a long single kinocilium and a bundle of shorter stereocilia (31). The immunofluorescent staining revealed App expression in both types of cilia at early time points in the larvae development (***Figures 2B-C***). Staining of 24 hpf larvae with glutamylated tubulin, highlighting the cilia basal body, clearly showed an App signal within the hair cells and close to the basal body (***Figures 2B, B’,B’’***). App expression became more distinct at 30 hpf (***Figure 2C***). Plots of the intensity profile of App (green) and acetylated tubulin (magenta) showed a punctate distribution of App throughout the kinocilium (***Figure 2D***), which supports that App localizes to the cilium membrane (***Figure 2E***). No signal was detected in the intensity profile in the absence of App puncta (***Figure 2F***), and the negative control (absence of primary antibody) was negative (***Supplementary file 2***). Moreover, a 3D-projection of the immunostaining similarly showed co-localization of App and acetylated tubulin (***Supplementary file 2A***). Together, these data show expression of App in cilia and ciliated cells of the otic vesicle and olfactory bulb and indicate that App is located at the cilia membrane.

### App localizes to cilia decorating the brain ventricle surface of zebrafish

As APP was previously shown to be expressed by the ependymal cells in rodents and in humans (32–34), we explored App expression by ependymal cells and App localization at their cilia in larvae and adult zebrafish (***Figure 3***). At 30 hpf, the brain ventricles are inflated and the differentiation of motile cilia in the most ventral and dorsal regions have just started but do not yet contribute to directional CSF flow (35). This facilitates whole-mount imaging and measurement of single cilium. Using the same combination of antibodies (anti-APP (Y188) and anti-acetylated tubulin) as above, we could detect App-positive puncta along the acetylated tubulin signal in most cilia (***Figures 3B,C***). To address if App localization to cilia is maintained into adulthood, we performed immunofluorescent staining on coronal sections of adult zebrafish brains using antibodies detecting App (Y188) and acetylated tubulin to label cilia. Our results showed that consistent to larvae, App was distributed to ependymal cilia in the adult brain. In contrast to larvae, ependymal cells in adult individuals were covered with multiple motile cilia. Cryosections of adult zebrafish brain revealed dense cilia tufts with App-positive staining at the apical side of the ependymal cells (***Figures 3E,F***). Furthermore, App was also expressed by ependymal cells, similarly to what has previously been described in rodents and humans (***Figure 3F***). Negative controls did not show any cilia-specific staining (***Supplementary file 3***).

**Figure 3.**
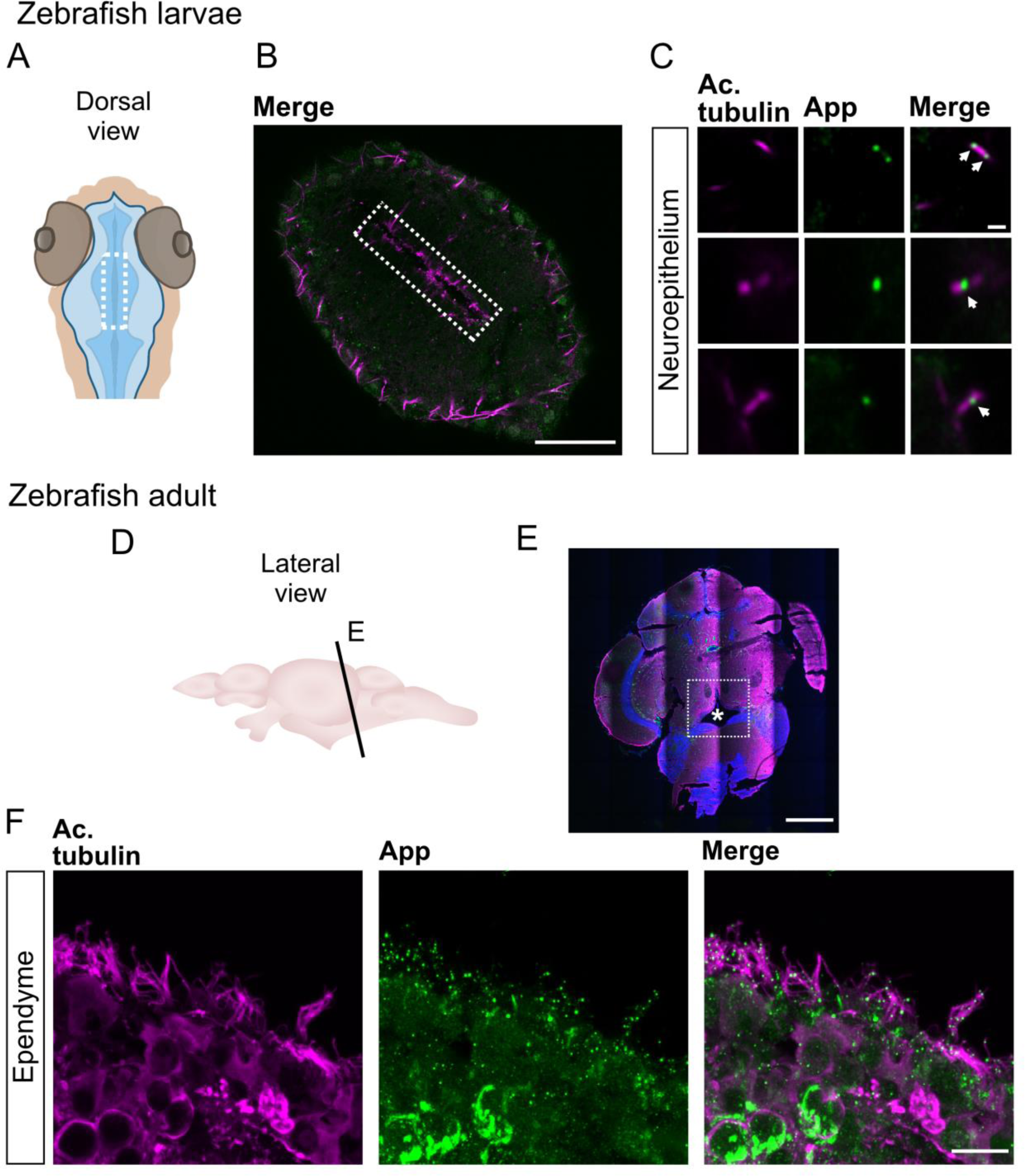
App is localized to the cilia membarne of neuropithelium and ependyme in larvae and adult zebrafish. App localize to the cilia decorating the ventricle of larvae and ependymal cells in adult zebrafish. (**A**) Schematic representations of head and ventricle morphology in 30 hpf zebrafish larvae, dorsal view. (**B**) Dorsal view of ventricle immunostained for App (green) and acetylated tubulin (magenta) in 30 hpf WT zebrafish larvae. (**C**) Close-up of cilia (magenta) and App (green). (**D**) Schematic outline of adult zebrafish brain, lateral view. (**E-F**) Coronal section of adult zebrafish brain and the central canal (see asterisk). Cell nuclei labeled with DAPI (blue), acetylated tubulin (magenta), App (green). (**F**) Ependymal motile cilia (magenta) of the central canal with App (green) accumulation along cilia. Magnification: (**B, E**)=10x, (**C, F**)=60x. Scale bar: (**B**)= 50µm, (**C**)= 1µm, (**E**)= 500µm, (**F**)= 10µm.

### Conserved localization of APP in ependymal cilia in mouse and human brains

APP is also localized to ependymal cilia in mice and humans. We performed immunostaining on mouse brain sections using two antibodies directed to APP, Y188 binding to the C-terminal intra-cellular domain and 22C11 detecting the E1 domain of the N-terminal region (***Figure 6C***), together with anti-acetylated tubulin. The ependymal motile cilia were easily localized in the third ventricle of the brain sagittal section (***Figures 4A,B***). Congruent with our results on adult zebrafish brains, we detected strong APP expression with both antibodies throughout the ependymal cells layer and punctate APP staining (Y188 see ***Figure 4C*** and 22C11 see ***Figure 4D***) overlapping with acetylated tubulin-positive cilia. Interestingly, APP expression by the choroid plexus cells was detectable (***Figure 4B***). Negative control for primary antibodies was performed and showed no or weak signal (***Supplementary file 4***).

**Figure 4.**
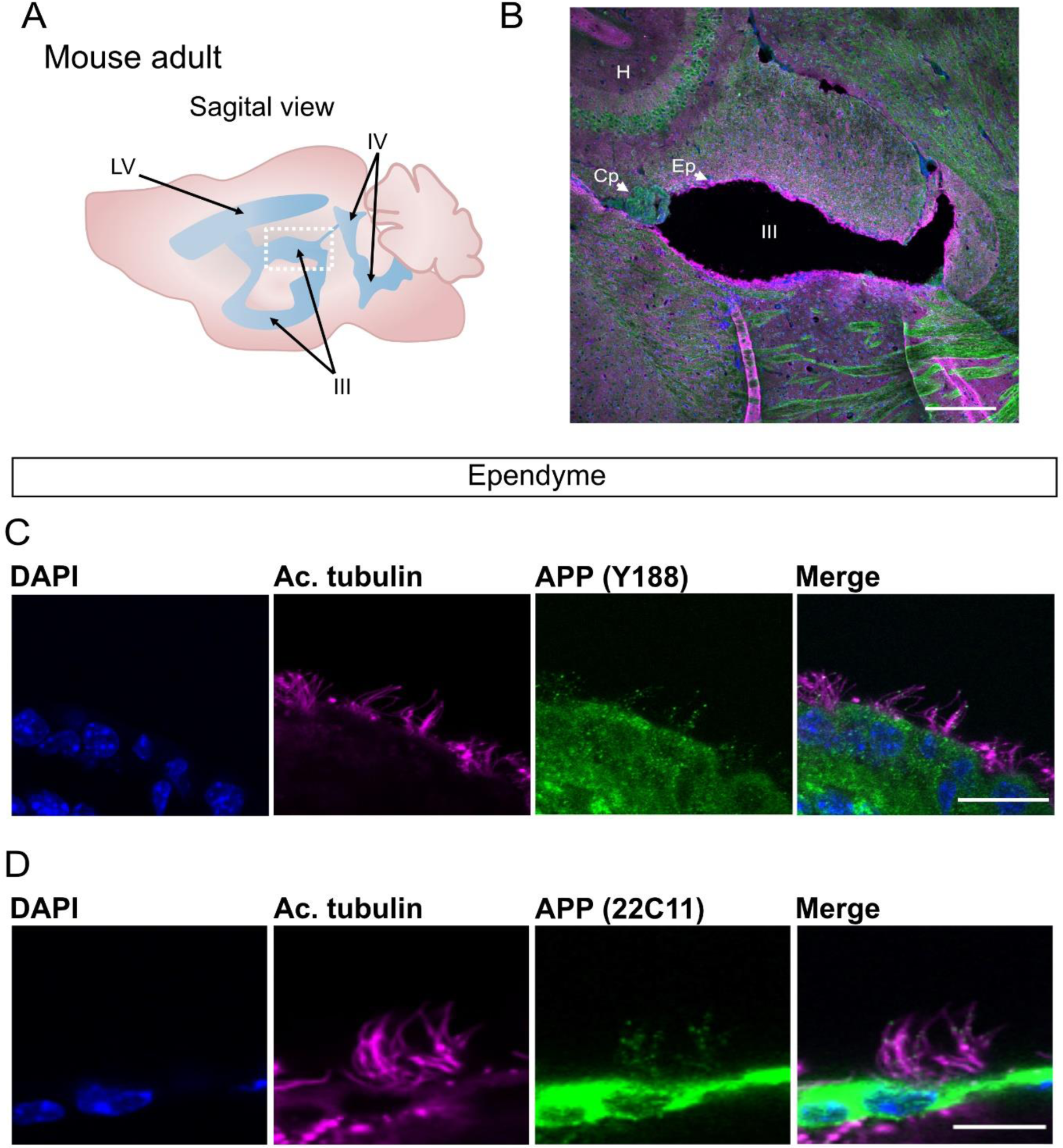
APP is localized to the ependymal cilia in adult mouse. APP is localized to the ependymal cilia in adult mouse. (**A**) Schematic representation of adult mouse brain ventricular system, sagittal view. (**B**) Overview of sagittal section from adult mouse brain and the third ventricle (see dotted square in (**A**)) for cell nuclei stained with DAPI (blue), acetylated tubulin (magenta), APP (green). (**C-D**) Close-up of ependymal cells and their cilia tufts (magenta) and APP expression with anti-APP Y188 antibody (**C**) and 22C11 antibody (**D**). LV= lateral ventricle, III= third ventricle, IV= fourth ventricle, H=hippocampus, Cp= choroid plexus, Ep= ependyme. Magnification: (**B**)=10x, (**C,D**)=60x. Scale bar: (**B**)= 200µm, (**C,D**)= 10µm.

In the human brain, acetylated tubulin staining allowed separation of cellular layers of the caudate nucleus and identification of acetylated tubulin-positive cilia of the ependymal cell layer lining the lateral ventricle (***Figures 5A,B,D***). However, while many ependymal cells had intact cilia, many were found broken and dislocated from their cell (***Supplementary file 5***).

**Figure 5.**
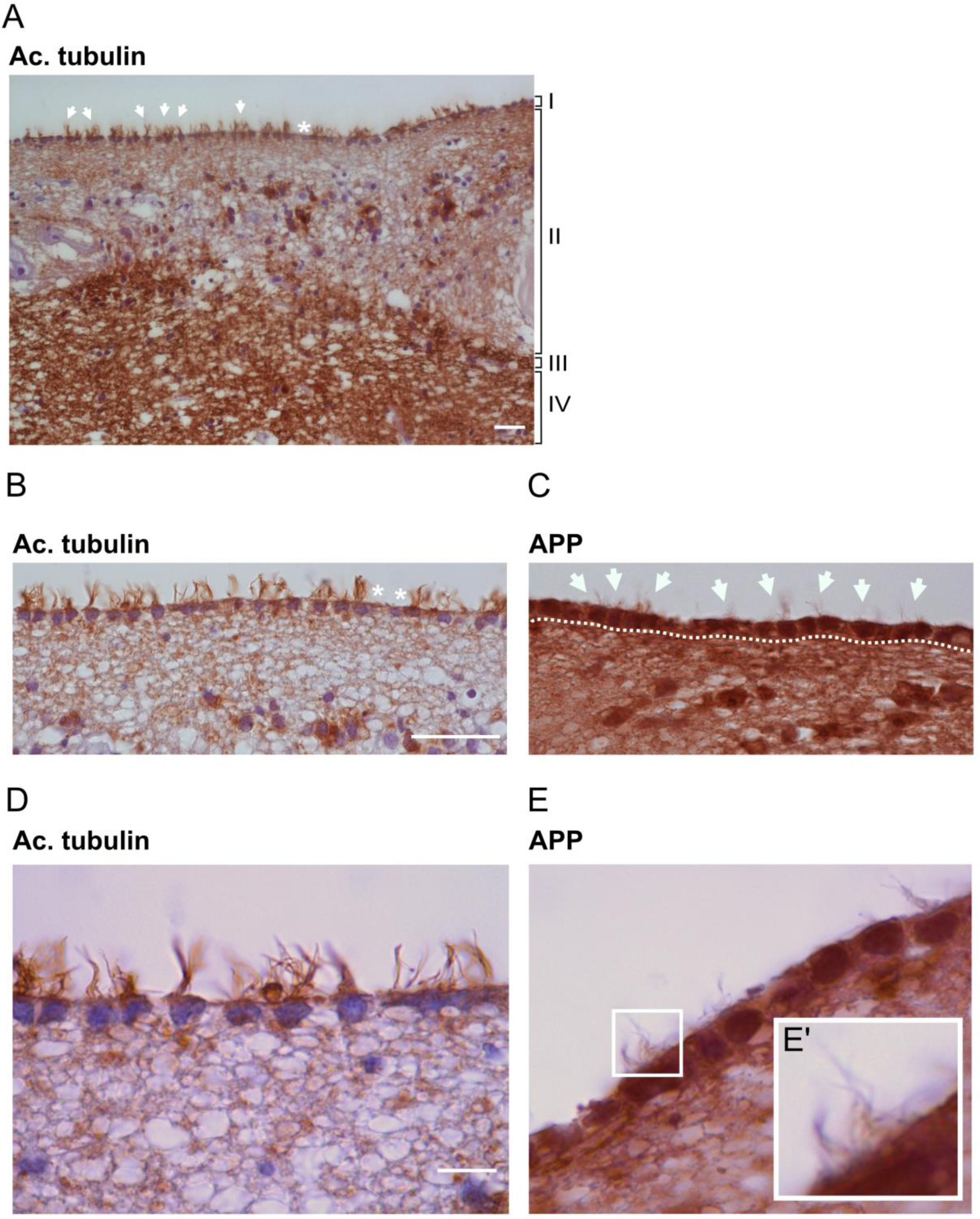
APP is localized to human brain ependymal cilia. APP is localized to human ependymal cilia. (**A**) Brightfield overview of a human brain section of the caudate nucleus immunostained with an anti-acetylated tubulin antibody reveals the different cellular layers (I-IV): (I) ependyme layer with motile cilia orienting towards the ventricle lumen, (II) cellular extensions connecting the ependymal cells, (III) cellular layer dense in astrocytes, (IV) brain parenchyma. (**B-E**) Higher magnifications of the ependymal layer show clear cilia (acetylated tubulin (**B,D**)) and APP (Y188 antibody (**C,E**)) accumulation within ependymal cells and along ependymal cilia. (**E’**). Arrows highlight ependymal cilia tufts in the ventricular lumen. White asterisks indicate broken or absent cilia. Dotted lines delimitate the ependymal cell layer. Magnification: (**A**)=20x, (**B-C**)= 40x, (**D-E**)= 100x. Scale bar: (**A**)= 5µm, (**B**)= 10µm, (**D**)= 2µm.

To address the presence of APP in ependymal cilia, brain serial sections of the caudate nucleus were incubated with horseradish peroxidase (HRP)-conjugated Y188 or A8717 antibodies, both recognizing the C-terminal domain of APP. Similarly to our results obtained in mouse and zebrafish brains, brightfield images confirmed strong APP expression in the ependymal cells and, upon higher magnification, in ependymal cilia (***Figures 5C,E*** , ***Supplementary file 5***). In contrast to zebrafish and mouse, APP in human ependymal cilia was evenly distributed and was not detected as puncta.

In summary, these results show that the expression of APP in the ependymal cells and their cilia are conserved between species as far apart as zebrafish, mice, and humans.

### Generation of appa and app double mutant zebrafish

In contrast to humans and mice, zebrafish have two *APP* orthologues, *appa* and *appb* (together designated *app*). Zebrafish with mutated *appb* gene was generated and described by our lab previously (7). However, to investigate the requirement of both App proteins in ciliogenesis, we used the CRISPR/Cas9 method to generate mutations in the zebrafish *appa* gene (***Figure 6A***). A mutation was identified in exon 2 (***Figure 6A***), and Sanger sequencing confirmed a frame shift mutation of 10 nucleotides (***Figure 6B***). The mutation resulted in a premature stop codon that is predicted to give rise to a protein truncation at amino acid 109 (***Figure 6C***). The *appa* mutant allele was outcrossed into the AB background until generation F4 and then bred with the *appb-/-* to generate the double mutant *appa*^-/-^*appb*^-/-^ zebrafish line. The *app* mutant zebrafish were healthy and fertile and did not show any gross morphological phenotypes. qPCR analysis of genes expression showed very low *appa* and *appb* mRNA levels in the double mutant fish line (***Figure 6D***). Western blot analysis using the Y188 and 22C11 antibodies with epitopes in the intracellular and extracellular domain, respectively, showed decreased protein levels in *app* double-mutant larvae (***Figure 6E***). Both antibodies are likely cross-reacting with Aplp2 since the epitope sequences are highly similar. These data show that the introduced mutation in *appa* resulted in a significant decrease of both transcription and translation of the Appa protein indicating that the mutation give rise to a loss-of-function mutation.

**Figure 6.**
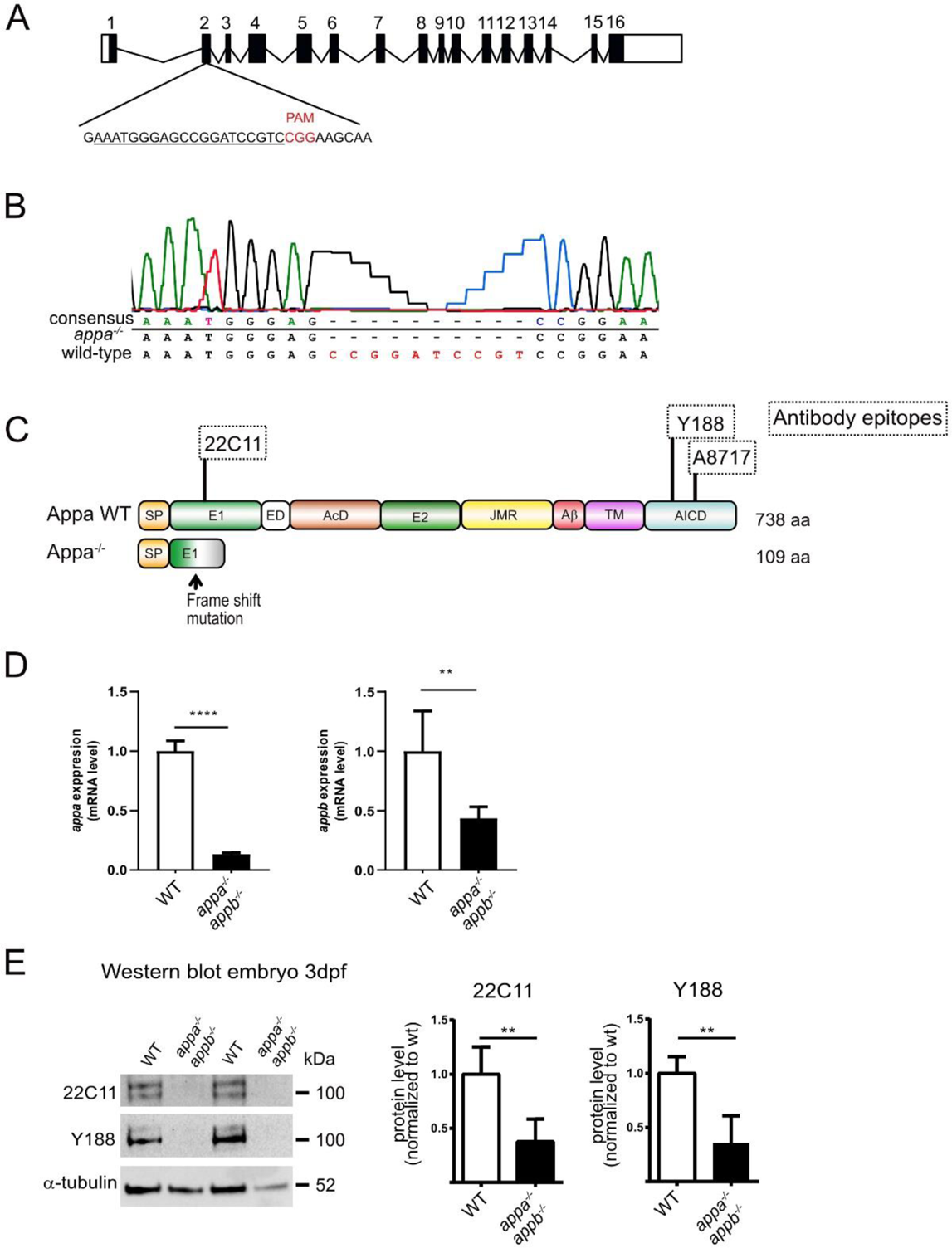
Generation of appa^-/-^ zebrafish. Generation of *appa^-/-^* and analysis of *appa^-/-^appb^-/-^* double mutant zebrafish. (**A**) Schematic outline of the *appa* gene with exons (black box) and UTR regions (white box). sgRNA used to target exon 2 with protospacer adjacent motif (PAM) in red and the sgRNA target sequence underlined. (**B**) Sanger sequencing chromatogram of exon 2 in wild-type and *appa^-/-^* zebrafish. (**C**) Schematic drawing of the wild-type Appa protein (738 aa) with epitopes of antibodies (dotted squares) used above and the hypothetical truncated Appa (109 aa) protein produced in *appa* mutant below. (**D**) qPCR quantification of *appa* and *appb* mRNA levels in wild-type and *appa^-/-^appb^-/-^* mutants at 24 hpf. (**E**) Western blot of 3 dpf whole larvae zebrafish with antibodies against 22C11 and App (Y188). Alpha-tubulin is used as loading control. Quantification of band intensity are shown relative to control. Data are reported as mean ± SD. ** ρ < 0.05, **** ρ < 0.001. qPCR n=5, WB n=3. SP= signal peptide, E1= extracellular domain, ED= extension domain, AcD= acidic domain, E2= extracellular domain 2, JMR= juxtamembrane region, Aβ= amyloid beta, TM= transmembrane, AICD= amyloid intracellular domain.

### Longer brain ventricle cilia in appa^-/-^appb^-/-^ larvae

The conserved distribution of APP in brain ventricle cilia prompted us to address the requirement of App during ciliogenesis. We measured the length of cilia in the midbrain ventricle detected by acetylated tubulin immunostaining signal in both *appa*^-/-^*appb*^-/-^ double mutants and wildtype larvae at 30 hpf. At this stage, the cilia delineating the dorsal and ventral parts of the diencephalic ventricles are not yet motile (35). A 3D-region of interest (ROI) was used to measure cilia length. The ROI was established from the dorsal part of the midbrain ventricle to the ventricular space at a depth of around 25 μm. To our surprise, we found that the ependymal cilia in the ROI were significantly longer in *appa*^-/-^*appb*^-/-^ mutants compared with wild-type larvae (***Figure 7***), which was confirmed by frequency distribution (***Supplementary file 7***).

**Figure 7.**
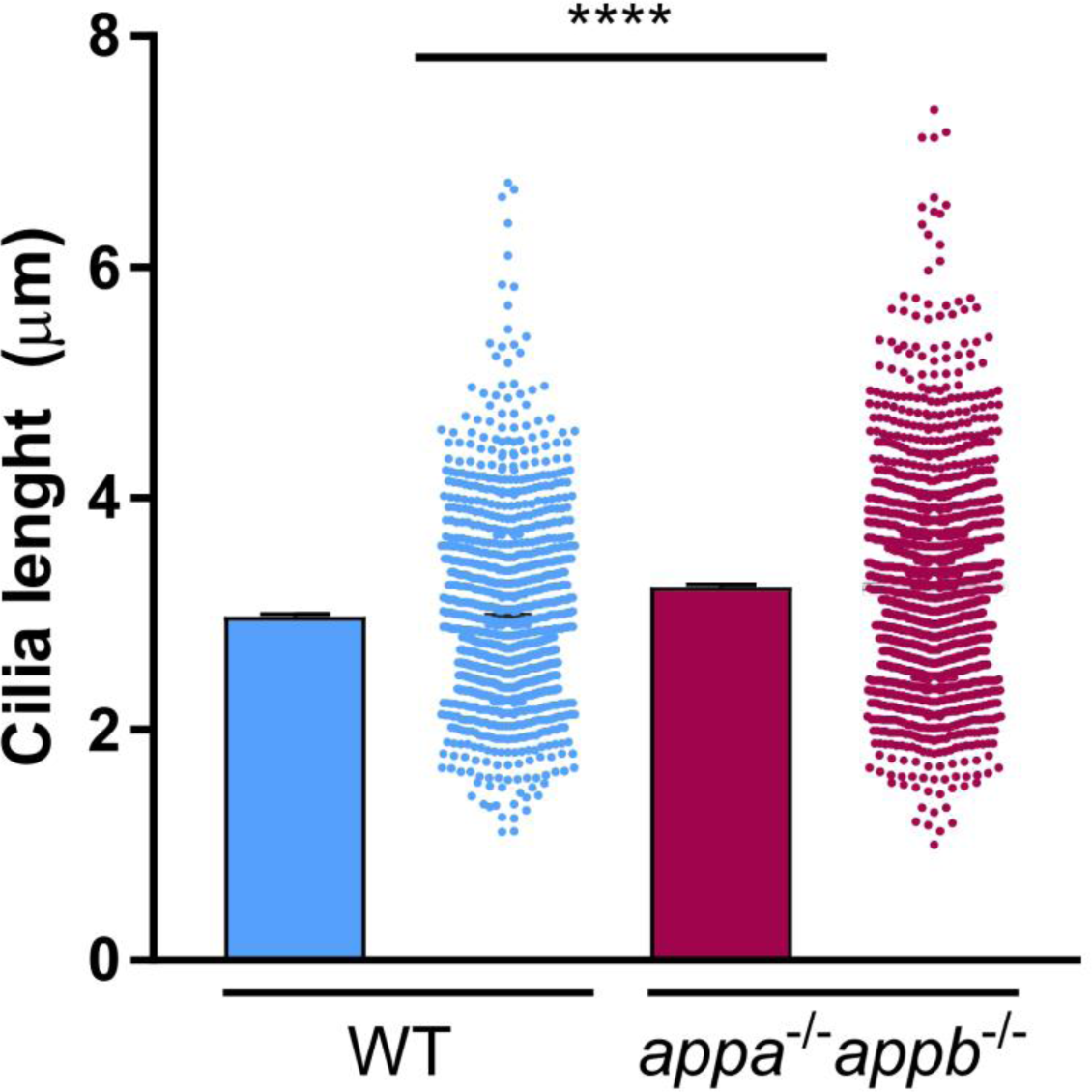
Longer cilia of brain ventricle neuroepithelium in *appa^-/-^appb^-/-^* larvae zebrafish. Longer cilia of dorsal brain ventricle neuroepithelium in *appa^-/-^appb^-/-^ larvae zebrafish.* At 30hpf, *appa^- /-^appb^-/-^* exhibit longer diencephalic/mesencephalic ventricle cilia than WT. Data are reported as mean ± SEM. **** ρ < 0.001. n=10 WT (1091 cilia), 16 *appa^-/-^appb^-/-^* (1511 cilia).

### Integrity of ependymal cilia axoneme and microtubule doublets in motile brain ependymal cilia in appa^-/-^appb^-/-^ mutant adult zebrafish

Emerging from the basal body is the axoneme, which forms the core of the cilium. First described in the early 1950s with electron microscopy, axonemes are composed of nine microtubule doublets at the periphery (9+0) (36). In some cilia, an additional central doublet is expressed (9+2), allowing cilia to generate and regulate movement (37, 38). This central microtubule doublet is found in motile ependymal cilia (9+2). To better characterize the ciliary ultrastructure of App-deficient zebrafish, we performed transmission electron microscopy (TEM) analysis of ependymal cells in adult zebrafish brains. TEM revealed a normal (9 + 2) axoneme in the cross-sections of ependymal cilia of WT (n=3) brain ventricle (***Figures 8A–D***). In *appa^-/-^appb^-/-^* zebrafish (n=4), ependymal cilia showed normal (9+2) axonemes (***Figures 8E– H***).

**Figure 8.**
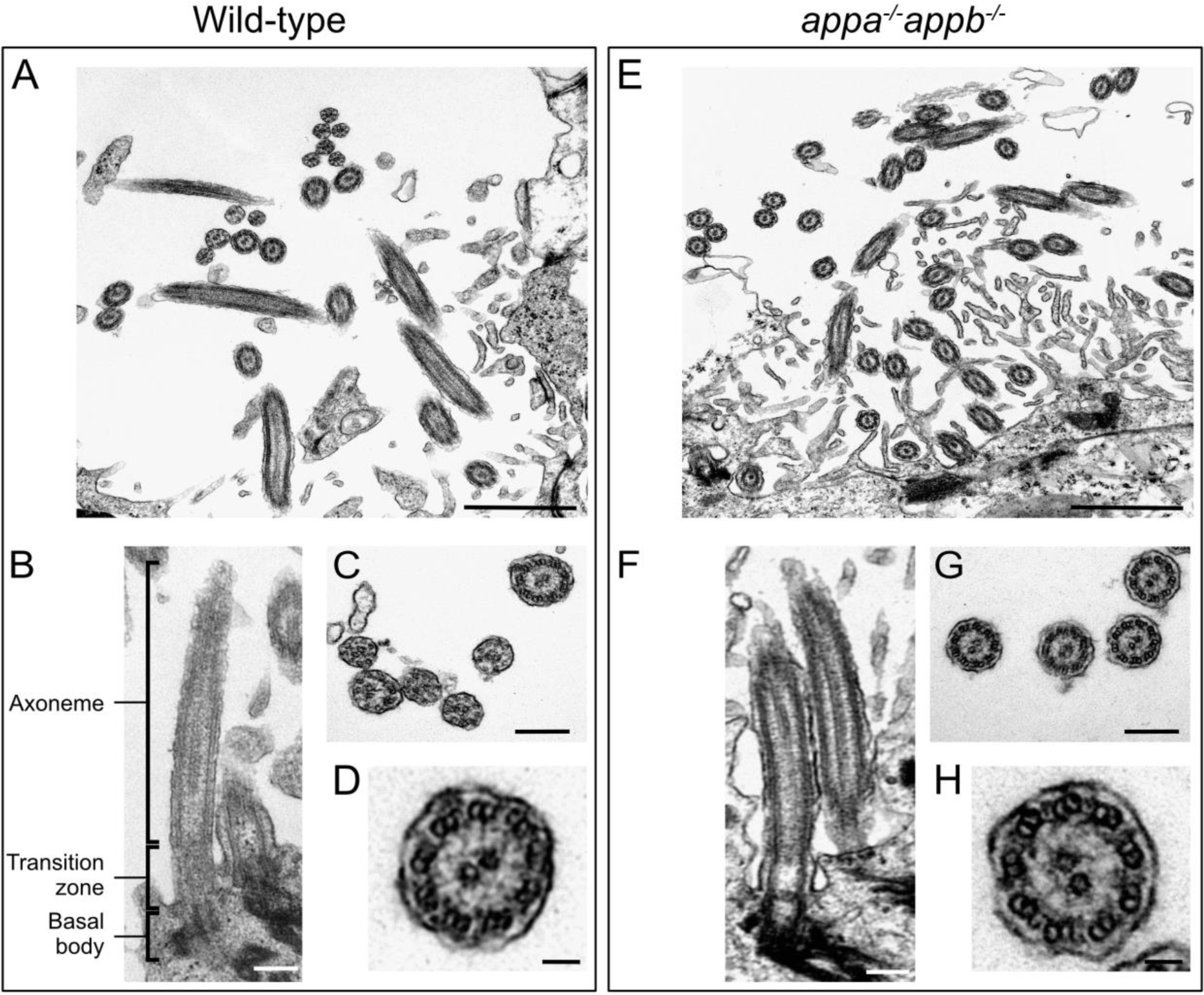
Structural integrity of ependymal cilia in WT and *appa^-/-^appb^-/-^* zebrafish. Transmission electron microscopy of adult zebrafish ependymal cilia of WT (**A**-**D**) and *appa^-/-^appb^-/-^* mutant (**E**-**H**) adult zebrafish. (**A**,**E**) Overview of ependymal cilia of the central canal. (**B**,**F**) Longitudinal view on the axoneme of the cilia composing its core, the transition zone including the ciliary pit between the cilia core and the cellular membrane and the basal body containing the cilia centrioles, highlighted with increased signal. In (**C**,**G**), cross-sections of cilia. (**D**-**H**) Zoom on cross-section of individual cilia showing (9+2) microtubule doublet organization. Scale bar: (**A**,**E**)= 1µm, (**B**-**C**, **F**-**G**)= 200 nm, (**D**,**H**)= 50 nm.

### The appa^-/-^appb^-/-^ double mutants exhibit smaller diencephalic ventricle

We then went on to address if defects in ependymal cilia affect brain ventricle formation. We analysed brain ventricle volume and area in 2dpf larvae (***Figure 9A***) and found significant reductions in both area and volume of the ventricular space in *appa^-/-^appb^-/-^* compared with wild-type (***Figure 9B***). These reductions were also observed when only the diencephalic ventricle was analysed (***Figure 9C***) and compared between both genotypes (***Figure 9D***). The gross morphology was next determined by measuring the length between specific points and areas of the ventricles: rostral to caudal, diencephalon ventricle sagittal length, amplitude and height (***Figure 9E***). However, no significant change was detected compared with wildtype larvae (***Figure 9F***). These results show that while the overall brain morphology of App mutants is maintained, their ventricles are smaller.

**Figure 9.**
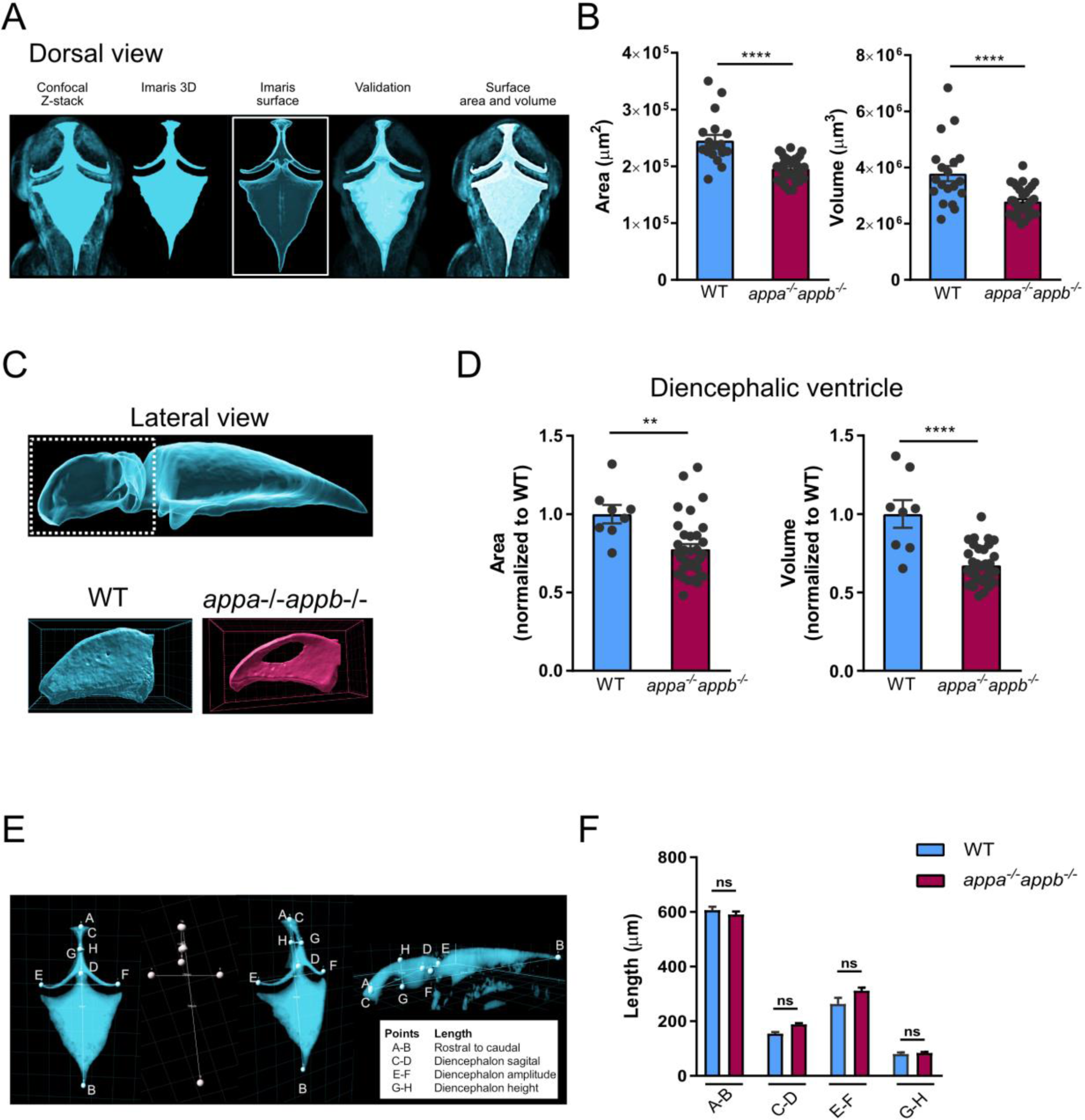
appa^-/-^appb^-/-^ larvae zebrafish exhibit smaller brain ventricle. The *appa^-/-^appb^-/-^* 2 dpf larvae zebrafish exhibit smaller brain ventricle. Dorsal 3D surface rending of confocal stacks taken from brain ventricles of dextran injected 2 dpf zebrafish larvae (**A**). Quantification of total ventricle surface area and volume show that both are decreased in *appa^-/-^appb^-/-^* larvae (**B**). Lateral 3D surface rending of confocal stacks from brain ventricles of dextran injected 2 dpf zebrafish larvae with close up on diencephalic ventricle (**C**). Quantification of surface area and volume of the diencephalic ventricle in WT and *appa^-/-^appb^-/-^* larvae (**D**). Measurement of gross ventricle morphology at 2dpf WT and *appa^-/-^appb^-/-^* larvae as the length (**E**). Distance between rostral to caudal, diencephalon ventricle sagittal length, amplitude and height show no significant difference in mutants (**F**). Data are reported as mean ± SEM. ** ρ < 0.01, **** ρ < 0.001. n: (**B**) WT=19, *appa^-/-^appb^-/-^* = 34, (**D**) WT=8, *appa^-/-^appb^-/-^* = 34, (**F**) WT=5, *appa^-/-^appb^-/-^* = 4.

### Cilia targeting motifs in App

Many proteins distributed to the cilium carry one or more cilia targeting sequences (CTS). The most common and well-studied are the VxP and AxxxQ motifs, both of which the requirement has been shown in transmembrane proteins including opsins (39–41) and somatostatin receptor 3 (SSTR3) (42, 43). The presence of App in cilia therefore made us investigate the presence of these motifs in App. Interestingly, we found several different CTS motifs with most localized to the mid- and C-terminal domain of the App protein (***Supplementary file 10***). Furthermore, most of these are in conserved regions and are thus shared between zebrafish, mouse and human (***Supplementary file 10***).

## Discussion

In this study, we show that App localizes to several different non-motile and motile cilia in zebrafish larvae including the stereo- and kinocilia of the otic vesicle, motile cilia of olfactory sensory neurons in the olfactory epithelium and cilia of the ependymal cells lining the brain ventricles. We also show an evolutionary conserved localization of APP to cilia of the ependymal cells lining the brain ventricles of adult zebrafish, mice and humans. As these results indicated a possible function of APP in ciliogenesis or cilia function, we used zebrafish lacking the two APP orthologues, Appa and Appb, and found longer ependymal cilia and smaller brain ventricles in larvae zebrafish. Thus, our results suggest that APP not only is distributed to cilia but also seems to have an important function in ciliogenesis and brain development.

### APP distribution within cilia

We used different antibodies to confirm the localization of App to the cilia. The punctate localisation of APP indicates that the protein is randomly distributed within the cilium similar to other membrane receptors such as SSTR3 and Smoothened (Smo) (44, 45). The distribution of APP within the plasma membrane varies between cell types, nonetheless a recent study suggested that APP clusters form groups of proteins within the plasma membrane (46). The similarity with the punctate pattern found here suggests that App may form clusters within the cilium, at least in zebrafish and mice. In contrast, we observed a continuous rather than punctate distribution of APP in human ependymal cilia. Whether the observed differences are due to sample preparation or variations in APP distribution between species remains to be addressed. Moreover, the accumulation of App at the root of the basal body, as observed in the olfactory sensory neurons and otic vesicle cilia in larvae zebrafish, correlates with the findings reported by Yang and Li on APP enrichment along ciliary rootlets (47).

The presence of APP within cilia raises the question of how APP is targeted to the cilium. The cilium membrane is continuous with the plasma membrane, yet it possesses a specific and conserved composition of proteins and lipids. This specification is considered to be established through an active transport of ciliary membrane proteins (48) that at least partly depends on specific ciliary transport sequences (CTSs) within the proteins (49). The presence of several such CTSs and their conservation between zebrafish, mice and human supports a motif-based transport of APP to cilia (***Supplementary file 9***). It will therefore be very interesting to address the extent to which these motifs are required for accumulation of APP at the root of the basal body and later distribution of APP out to the cilium.

### App in brain ventricles

APP expression by the ependymal cells was first reported in rodents and humans in the late 1980s and early 1990s (32, 33, 50, 51). In line with these findings, our results not only confirm the expression of App in adult zebrafish ependymal cells, but in addition, show that APP localizes to ependymal motile cilia in vertebrates as far apart as zebrafish, mice, and humans. Our finding that loss of App results in morphologically abnormal ependymal cilia suggests a role of App in ciliogenesis. However, the *appa^-/-^appb^-/-^* mutants gave rise to fertile adults without major phenotypic changes associated with cilia defects, such as curved body and hydrocephalus (52–55). In line with our findings, Olstad *et al*. recently reported that such phenotypes mainly associate with primary cilia defects, while changes in motile cilia were more likely to result in ventricle duct occlusion (56). During early development, movement of cilia is a major factor maintaining CSF flow within ventricles. Consequently, the cilia-driven flow is crucial to form and maintain a proper brain ventricular system, as zebrafish, clawed frog and mouse ciliary mutants display ventricular defects (56). It is thus likely that the defective ventricle expansion observed in the *appa^-/-^appb^-/-^* mutant larvae may result from changes in of motile cilia. Although we did not observe a lack of diffusion between ventricles indicating duct occlusion, our data suggest that App may be required in motile cilia to promote flow of CSF needed for ventricle formation during early development (56). The structural changes in cilia observed in adult *app* mutant zebrafish are similar to those observed in the cilia and flagella-associated protein (*CFAP43*) mutant mice with a normal pressure hydrocephalus-like phenotype (57). It will thus be interesting to examine the extent to which cilia movement and CSF flow change when altering App levels.

### The function of App in ependymal cilia

Our findings raise several questions regarding the role of APP in cilia and to which level cellular processes associated with APP may be mediated through cilia. For example, the multi-ciliated ependymal cell layer covering the brain ventricles is important for neurogenesis, both by regulating the number of neural stem cells in the neurogenic niches of the subventricular zone (51) and by facilitating the migration of new-born neuronal cells through cilia-regulated fluid dynamics (58). Interestingly, APP is also reported to regulate neurogenesis (59–61) and to promote neuronal migration (9). Our understanding of APP-mediated processes continues to increase, but the mechanisms by which these processes are orchestrated are yet not fully understood. Therefore, the obvious overlap between functions mediated by ependymal cells and APP makes it tempting to speculate that APP might be important or at least partly required in ependymal cells to mediate cell migration and proliferation.

The length of cilia can be modified both by changes in the structural proteins involved in microtubule assembly but also depends on the cell proliferation and differentiation status, where proliferating cells generally have shorter cilia than growth arrested cells. As an example, modulation of several cell cycle-related kinases could alter ciliary length (62). This is true for primary cilia but to what extent this is valid for other types of cilia is not yet described. The mechanisms by which App contributes to regulate cilia length is beyond the scope of the present study but could potentially involve its role in cell differentiation (6).

It is intriguing to think of APP in the cilium as a receptor that senses signalling molecules and metabolites transported through the ventricles by CSF. The hypothesis of APP acting as a receptor is supported by its similarities with type I membrane receptors and by the fact that the list of potential APP ligands continues to grow (review by (6)). Therefore, it is tempting to speculate that APP localized on the cilia interacts with CSF-circulating ligands, *e.g.*, Aβ peptides, growth factors, and hormones (6, 63), to mediate CSF-derived signalling.

### Long-term defects clearance

Beyond the impact of App on ciliogenesis during development, it is intriguing to speculate on the long-term effects of impaired near-wall CSF propulsion. This movement is thought to play an important role in removal of waste products from the brain parenchyma (64). Thus, it is likely that subtle changes in the coordinated beating of cilia may contribute to altered regional CSF flow that impairs clearance and hence contributes to a slow build-up of waste products over time. In support are findings that individuals with Down syndrome, expressing approximately 50% higher levels of APP, have changed CSF flow in the lateral ventricles (65). Although the morphology of ependymal cilia of DS brains are unknown, *in vitro* cell cultures show decreased primary cilia length (26). Investigations addressing cilia morphology and function in the adult zebrafish brain lacking App are ongoing in our lab; however, it will be equally important to perform these experiments in *app*-knockout mice, as well as in mice overexpressing APP, which results in altered post-translational processing of the protein.

### APP fragmentation and CSF biomarkers

The presence of APP in ependymal cells and their cilia also raises the question regarding their contribution to APP-derived fragments found in CSF. As at least some secretases needed for APP processing are present in cilia (66), it is likely that the fragments detected in CSF not only originates from the brain parenchyma but also from APP being processed within the ependymal cells and the protruding cilia. The release of APP from ependymal cells could be mediated through the release of extracellular cleavage products or by budding extracellular vesicles and ectosomes. The latter process was in a recent study described as a common mechanism by which proteins are cleared from cilia instead of recycling by retrograde transport (67). APP-containing vesicles are released into the CSF (68) and in a recent report, such microvesicles were found to have lower levels of APP in AD patients compared to healthy individuals (69). The impact of ependymal integrity and the contribution of cilia-mediated APP release need further studies but could potentially contribute to our interpretation of biomarkers used to assess disease progression. Interestingly, a well-established feature of normal pressure hydrocephalus, where ciliary function is impaired (70), is decreased CSF levels of soluble APP and Aβ, which are restored upon successful shunt treatment of the condition (71–73).

### App in otic vesicle and olfactory epithelium

#### Olfactory sensory neurons cilia

While others have shown that APP and its processing machinery are expressed in the olfactory epithelium and bulb in cultured mouse cells (74), we here report App localization also to the olfactory cilia in larvae zebrafish. Motile cilia of the olfactory sensory neurons (OSNs) in zebrafish are essential to generate liquid flow in the nose pit to detect odorant molecules (75). In zebrafish, the olfactory epithelium can be divided into three categories of OSN, *i.e.*, ciliated, microvillus and crypt OSNs (review by (76)). Each OSN expresses distinct classes of receptors and sensing molecules and has a specific axonal pathway from the olfactory bulb leading towards higher olfactory centres in either the telencephalon or the optic tectum. If APP is present in cilia of all OSN or only a subset, needs to be confirmed. However, the presence of APP in the olfactory cilia could potentially give clues on corresponding pathways and insights into the mechanisms resulting in olfactory deficiencies in AD mouse models and neurodegenerative disease (77).

#### Otic vesicle cilia

Hearing is a major sensory input in vertebrates, which is known to decrease with aging. Although the relationship between APP and hearing is less studied than many other areas, there are a few reports pointing to the loss of hearing associated with APP or its cleavage product Aβ (78–80). Our data, showing the presence of App in cilia mediating hearing, open up the possibility that nervous system-related changes in hearing may not only be due to defects in the brain regions receiving input from the auditory organ but also due to direct effects on the cilia. However, the function of App in the auditory system needs further investigation.

### Conclusion

Altogether, our data show the presence of App in motile and non-motile cilia of the otic vesicle, olfactory pit and ependymal cells lining the brain ventricles. We also report a conserved distribution, at least in the ependymal cilia, across vertebrates and that App is required for proper ciliogenesis and brain ventricle formation. The evolutionary conserved CTSs of APP and its expression throughout development and aging suggest a central role of APP within the ependyme. Further studies are required to fully understand the impact of App in cilia in our olfactory and auditory organs and to which extent defects in ependymal cell integrity and ciliation contribute to APP-related developmental processes and disease progression.

## Materiel and Methods

### Animal care and ethics statement

The zebrafish (*Danio rerio*) facilities and maintenance were approved and follow the guidelines of the Swedish National Board for Laboratory Animals. Experimental procedures were approved by the by the ethical committee in Gothenburg. Zebrafish were maintained in Aquatic Housing Systems (Aquaneering, San Diego, CA) at 28.5 °C, under a 14:10 hour (h) light:dark cycle at the Institute of Neuroscience and Physiology, University of Gothenburg. Fish were fed twice daily a diet of live-hatched brine shrimps and Gemma fish food (Skretting, Amersfoort, Netherlands). System water was created using reverse osmosis water kept at a pH of 7.2-7.6 with NaHCO3 and coral sand and salt (Instant Ocean, Blacksburg, VA) to maintain the conductivity at 600μS. Breeding of fish was carried out in 1-2 L breeding tanks and embryos were collected in embryo medium (EM) (1.0mM MgSO4, 0.15mM KH2PO4, 0.042mM Na2HPO4, 1mM CaCl2, 0.5mM KCl, 15mM NaCl, 0.7mM NaHCO3) and raised in a dark incubator at 28.5 °C (81).

The following fish lines were used in the present project; AB fish from the Zebrafish international resource centre (ZIRC) or was used for outbreeding and as wild-type background, *appb^26_2-/-^* (7) and *appa^-/-^* as described below.

### Mutagenesis using the CRISPR/Cas9 system

Genetic mutations in the *appa* gene were introduced using the CRISPR/Cas9 system as previously described (82). Briefly, gRNAs were generated with a target-specific DNA oligonucleotide (Integrated DNA Technologies, Leuven, Belgium) containing a T7 promoter sequence in the 5’-end and a ‘generic’ DNA oligonucleotide for the guide RNA. The two oligonucleotides were annealed and extended with Platinum Taq DNA polymerase (ThermoFisher, Waltham, MA), in a final concentration of 1x buffer, 0.25mM dNTP, 0.5μM of each oligonucleotide and 0.04U/ul Taq with one cycle at the following temperatures (98°C 2 min; 50°C 10 min, 72°C 10 min). The resulting product was analyzed on a 2.5% agarose (Roche, Basel, Switzerland) gel to confirm a single fragment of 120 basepairs (bp) and used to transcribe RNA. *In vitro* transcription was performed with the T7 Quick High Yield RNA Synthesis kit (New England Biolabs, Ipswich, MA) and incubated at 37°C for 16 h. DNA template was removed with RNase Free DNase at 37°C for 15 min. After purification with the RNA clean & concentrator-5 (Zymo Research, Irvine, CA), gRNA was analyzed on a 2.5% agarose gel for integrity and diluted to 250μg/μl with RNase free water and stored at -80°C. Cas9 protein was diluted to 500nM in Hepes (20mM HEPES, pH7.5; 150mM KCl) and stored at -80°C. Embryos were co-injected with 50 pg gRNA and 300 pg Cas9 protein at the one to two cell stage using a microinjector apparatus FemtoJet^®^ express (Eppendorf AG, Hamburg, Germany). Injected embryos were screened for gRNA activity using the T7 endonuclease assay (New England Biolabs, Ipswich, MA). Ten embryos from each gRNA injection were pooled at 48 hpf and genomic DNA extracted with 50mM NaOH at 95°C for 30 min. M13- and PIG- tailed primers (IDT, Leuven, Belgium) were used to amplify a region surrounding the mutated site of each locus using 1x buffer, 2.5mM MgCl2, 0.2mM dNTP, 0.2μM primers, 1U Taq polymerase (Promega, Fitchburg, WI). The polymerase chain reaction (PCR) was purified on an 1% agarose gel with the QIAquick Gel Extraction Kit (Quiagen, Hilden, Germany) and then 200 ng of the purified PCR product was dissociated and reannealed (95°C for 5min, 95-85°C at -2°C /s, 85-25°C at 0.1°C /s) in a reaction containing 1x NEB buffer 2 (New England Biolabs, Ipswich, MA) and then digested with 5U T7 endonuclease I (New England Biolabs, Ipswich, MA) for one hour at 37°C. Fragments were analyzed on a 2% agarose gel. The remaining embryos were raised to adulthood and outcrossed with AB wild-type fish. Sixteen embryos from each outcrossed pair were screened for mutations in the F1 generation using a three-primer fluorescence PCR method. A 300-450 bp region surrounding the target site was amplified using forward primers linked with a M13 sequence and a PIG-tailed reverse primer in combination with a generic M13-FAM primer. The *appa^C21_16^* mutants, refer to as *appa^-/-^*, carry a deletion of -10 bp in exon 2. Sanger sequencing with BigDye™ Terminator v1.1 Cycle Sequencing Kit (Applied Biosystems™, Waltham, MA) on an ABI3130xl sequencer (SeqGen Inc, Los Angeles, CA) revealed a deletion of ten nucleotides in exon 2 that likely introduce a frameshift mutations. Heterozygous mutant carriers were raised and subsequently outcrossed into the wild-type AB fish line until generation F4. Outcrossed adults were genotyped using M13-FAM primers and PCR reactions diluted in HiDi^TM^ formamide (Applied Biosystems™, Waltham, MA) with ROX^TM^500 dye size ladder (ThermoFisher, Waltham, MA) and analyzed for amplified fragment length polymorphism (AFLP) on an ABI3130xl sequencer. Offspring from heterozygous F4 inbreeds were inbred to generate homozygous wild-type and mutant lines. Generation of *appa^-/-^appb^-/-^* double mutants were obtain from mating single mutant *appa^-/-^* with single mutant *appb^-/-^*.

### Protein sequence alignment

Sequences of APP were obtained from the UniProt database (83) and aligned with ClustalW using MegAlign Pro v17.2.1 (DNAstar, Inc., Madison, WI) The following sequences were used; *Homo sapiens* APP751 (P05067-8), *Mus musculus* APP751 (P12023-3), *Danio rerio* Appa738 (Q90W28), Appb751 (B0V0E5). Amino acids conserved across all species were marked with bright blue background.

### Whole-mount fluorescent in situ hybridization

To detect *appa* and *appb* mRNA expression pattern in zebrafish larvae, fluorescent *in situ* hybridization was performed. Antisense digoxigenin-labeled *appa* and *appb* RNA probes used are described previously (84). Zebrafish embryos were staged according to Kimmel *et al*. to the hours post-fertilization (hpf) (85) and manually dechorionated with forceps (Dumont, Montignez, Switzerland). A treatment with 0.003% PTU (1- phenyl-2-thiourea) (Sigma, St. Louis, MO) was performed around 23hpf stage to prevent pigmentation. Fluorescent *in situ* hybridization was performed as described by Lauter *et al*. (86). Briefly, zebrafish larvae were euthanized in 0.2mg/ml ethyl 3-aminobenzoate methanesulfonate (tricaine) (MS-222, Sigma, St. Louis, MO) (81) and fixed at 30 hpf in 4% paraformaldehyde (PFA) (Sigma, St. Louis, MO) for 24h at 4°C. Embryos were washed in phosphate-buffered saline (PBS) with 0.1%Tween-20 (PBST) and dehydrate into increasing methanol (MeOH) gradients from 25 to 100%. Embryos were incubated in 2% hydrogen peroxide (H2O2) for 20 min, then gradually rehydrated with decreasing MeOH gradients. Embryos were incubated in 10μg/ml proteinase K (in 10mM Tris-HCl pH 8.0, 1.0 mM EDTA) for 10 min at room temperature (RT). The reaction was stopped with 2 mg/ml glycine in PBST and then the embryos were postfix in 4.0% PFA for 20 min. PBST washes were performed before incubation in prehybridization buffer (HB; 50% deionized formamide, 5x saline-sodium citrate (SSC) (3M NaCl, 300 mM tri-sodium citrate, pH 7.0), 5 mg/ml torula RNA (Sigma, St. Louis, MO), 50 μg/ml heparin sodium salt and 0.1% Tween- 20). Embryos were pre-hybridized at 70°C for 1h. Then, hybridization was done with selectively 50 ng of DIG-labelled *appa* or *appb* RNA in HB with 5% dextran sulfate (Sigma, St. Louis, MO) at 70 °C overnight. The next day, embryos were washed in warm SSC with 0.1% Tween-20 followed by PBST only. After that, a 1h-blocking incubation at RT in PBST with 8% goat serum (Sigma, St. Louis, MO) was performed. For the antibody treatment, a sheep-anti-digoxigenin-peroxidase (POD)-Fab fragments antibody (1:500 in blocking solution) (Roche, Basel, Switzerland) was used and embryos were incubated in the dark overnight at 4°C, without agitation. To remove excess antibody, embryos were then washed in PBST at RT in gentle agitation. To amplify the signal, tyramide signal amplification (TSA) was used by combining 5-carboxyfluorescein succinimidyl ester (Molecular Probes, Eugene, OR) with tyramine hydrochlorine (Sigma, St. Louis, MO) at a 1.1:1 respective equimolar ratio. Vanillin (0.45mg/mL) (Sigma, St. Louis, MO) was used as a POD accelerator and diluted in borate buffer pH 8.5. Embryos were incubated with the TSA and POD accelerator reaction in the dark without agitation for 15 min at RT. To stop the TSA reaction, embryos were washed in PBST and then incubate in 100 mM glycine-HCl pH 2.0 to inactivate the POD reaction followed by additional PBST washing. To avoid shrinkage, embryos were then incubated in an increasing glycerol gradient (in PBST, 40mM NaHCO3). Whole embryos were mounted on glass bottom 35 mm Petri dish (Cellvis, Mountain View, CA) in 1% low-melting agarose (Sigma, St. Louis, MO). Samples were imaged as stacks using inverted Nikon A1 confocal system (Nikon Instruments, Melville, NY) using a 20x objective (Plan-Apochromat 20x/0,75) and 40x water-immersion objective (Apochromat LWD 40x/1,15). Image processing was done using ImageJ FIJI software (NIH, Bethesda, MD).

### Immunofluorescence

#### Zebrafish larvae

To detect protein expression, immunofluorescence experiments were performed in whole-mount AB zebrafish larvae. A treatment with 0.003% PTU was performed around 23hpf stage to prevent pigmentation. Then freshly euthanized embryos were fixed at 30 hpf for 2h in 4% PFA at RT on slow agitation. After fixation, embryos were washed with PBS with 0.5% Triton-X (PBTx) at RT. Followed up by incubation in blocking solution (5% goat serum donor herd (GS) (Sigma, St. Louis, MO), 2% bovine serum albumin (BSA) (Sigma, St. Louis, MO) , 1% DMSO (Sigma, St. Louis, MO) and 0.5% PBTx) for 3h at RT. The larvae were then incubated overnight at 4°C on slow agitation with the desired primary antibodies in blocking solution: mouse IgG2b anti-acetylated tubulin monoclonal antibody (1:1000) (Sigma, St. Louis, MO), recombinant rabbit anti-amyloid precursor protein monoclonal antibody Y188 (1:500) (Abcam, Cambridge, United Kingdom), and/or mouse anti-glutamylated tubulin monoclonal antibody (1:1000) (Adipogen, San Diego, CA). The zebrafish larvae used for negative control were incubated in blocking solution only. The next day, embryos were washed (5x 45min) with PBSTx at RT and incubated in dark with the specific secondary antibodies overnight at 4°C, in blocking solution: goat anti-mouse IgG2b Alexa Fluor-647 (1:1000) (Invitrogen Thermo Fisher, Waltham, MA) and goat anti-rabbit IgG Alexa Fluor-488 (1:1000) (Invitrogen Thermo Fisher, Waltham, MA), or goat anti-mouse IgG1 Alexa Fluor-568 (1:1000) (Invitrogen Thermo Fisher, Waltham, MA). The zebrafish larvae used for negative control were also incubated with the former secondary antibodies. The larvae were then washed with PBTx at RT and incubated for 15 min with DAPI (1:1000) (ThermoFisher, Waltham, MA) to stain the nuclei in PBS at RT before the final washes. Stained larvae were mounted in 1% low-melting point agarose, on glass bottom 35 mm Petri dish.

#### Adult zebrafish and mouse brains

Brains from adult zebrafish (AB, 2 year-old) and mouse (C57Bl6/n, 8-9 week-old). Brains were fixed in 4% PFA in PBS overnight at 4°C and then washed and immersed in 30% sucrose solution in PBS, after which they were frozen in OCT cryomount (Histolab, Askim, Sweden). Coronal or sagittal cryosections from adult zebrafish (25 μm) and mouse brains (16 μm) slices were stored at -80°C prior to use. Sections were air dried for 15 min at RT then rehydrated in PBS. Slices were permeabilized in 0.1% PBTx for 10 min at RT and washed 3x in PBS for 15min each. A 0.1% Sudan Black B (SBB) (Sigma, St. Louis, MO) in 70% EtOH treatment was performed for 20 min at RT. Slides were then washed in PBS for 3x5 min. The slides were then incubated in blocking solution of 2% GS in PBS at RT for 1h, followed by the incubation with the primary antibodies in 2% BSA at 4°C overnight: mouse IgG2b anti-acetylated tubulin monoclonal antibody (1:1000), recombinant rabbit anti-amyloid precursor protein monoclonal antibody (Y188) (1:500) or mouse anti-amyloid precursor protein A4 antibody (clone 22C11) (1:500) (Merck Millipore, Burlington, MA), or rabbit IgG (1:500) (Abcam, Cambridge, United Kingdom) and/or with blocking solution only for negative controls. The next day, slides were wash 3x in PBS for 15min each and incubated with the secondary antibody in 2% BSA at RT for 3.5h combined with DAPI (1:1000): goat anti-mouse IgG2b Alexa Fluor-647 (1:1000) and/or goat-anti rabbit Alexa Fluor-488 (1:1000) and/or goat anti-mouse IgG1 Alexa Fluor-488 (1:1000) (ThermoFisher, Waltham, MA) and/or goat anti-mouse IgG1 Alexa Fluor-568 (1:1000). The slides were then washed 3x15 min in PBS and shortly rinsed in ddH2O to remove any residual salts. The slides were covered with coverslips using ProLong gold antifade mounting medium (Invitrogen Thermo Fisher, Waltham, MA).

Samples were imaged using Zeiss LSM710 inverted confocal microscope (Carl-Zeiss, Jena, Germany) using 40x water immersion objective (Plan-Apochromat 40x/1.0) and a 63x oil-immersion objective (Plan-Apochromat 40x/1.0) or with Zeiss LSM880 Airyscan inverted confocal microscope (Carl-Zeiss, Jena, Germany) using 40x water immersion objective (LCD- Apochromat 40x/1.0) and 63x oil-immersion objective (Plan-Apochromat 63x/1.4). Image processing and intensity profiles were performed with ImageJ FIJI program.

#### Human brain sections immunofluorescent staining

Neurologically normal human post-mortem control tissue was obtained from Queen Square Brain Bank for Neurological Studies. Paraffin-embedded sections were cut from caudate nucleus brain region, which contains ependymal lining containing cilia. Sections were dewaxed in three changes of xylene and rehydrated using graded alcohols. Endogenous peroxidase activity was blocked using 0.3% H_2_O_2_ in MeOH for 10 min followed by pressure cooker pre-treatment for 10 min in citrate buffer, pH 6.0. Non-specific binding was blocked using 10% non-fat dried milk (Sigma-Aldrich, St. Louis, MO) in Tris buffered saline-Tween (TBS-Tween) before incubating with either anti-acetylated tubulin (1:1000) or anti-APP (1:500) antibodies at RT for 1 h. A biotinylated mouse anti-rabbit IgG antibody (1:200) (Agilent DAKO, Glostrup, Denmark) was added for a 30 min incubation with the sections at RT followed by avidin-biotin complex (Vector Laboratories, Burlingame, CA). Coloration was developed with diaminobenzidine (Sigma-Aldrich, St. Louis, MO) activated with H_2_O_2_ (87).

#### Protein extraction from whole zebrafish larvae and western blotting

Protein was extracted from 3dpf double *appa^-/-^appb^-/-^* mutant whole larvae (60 larvae per n, n=3) to confirm loss of protein. Larvae were euthanized, deyolked with ice-cold PBS and snap frozen in liquid nitrogen prior to use and stored at -80°C. Samples were homogenized in an ice-cold lysis buffer (10 mM Tris-HCl pH 8.0, 2% sodium deoxycholate, 2% SDS, 1 mM EDTA, 0.5 M NaCl, 15% glycerol) supplemented with protease inhibitors cocktail (Roche, Basel, Switzerland) and using glass tissue grinder, on ice. Samples were then incubated 20 min on ice, sonicated for 10 min on max level and centrifuged at 10,000 x g at 4°C. Supernatants were collected and kept on ice and protein concentration measured with a BCA Protein Assay Kit (ThermoFisher, Waltham, MA) and samples stored at -80°C. Proteins samples (40-60ug) were then diluted in a denaturing lysis buffer (1X NuPAGE^®^ LDS Sample Buffer (ThermoFisher, Waltham, MA), 0.05M DTT (Sigma-Aldrich, St. Louis, MO), lysis buffer completed with protease inhibitors) and then boiled for 5 min at 95°C. Proteins were then separated on a NuPAGE^®^ NOVEX^®^ Bis-TRIS pre-cast gel (Invitrogen Thermo Fisher, Waltham, MA) and transferred onto a 0.2 μm nitrocellulose membrane (GE Healthcare, Chicago, IL). The membrane was incubated in a blocking solution (5% milk) for 2h at RT and then immunoblotted with the desired primary antibodies overnight at 4°C: rabbit anti-amyloid precursor protein monoclonal antibody (Y188) (1:2000) or mouse anti-amyloid precursor protein A4 antibody (clone 22C11) (1:5000) and with a loading concentration control mouse anti-GAPDH-HRP conjugated (1:20000) (Novus Biologicals, Centennial, CO) or mouse anti-α-tubulin monoclonal (1:10000) (Sigma, St. Louis, MO). The membrane was then washed in TBS-Tween 3x 10min at RT and incubated with the secondary antibodies anti-rabbit-HRP (1:5000) (Cell Signaling, Danvers, MA) for 1h at RT. The membrane was washed 3x10min in TBS-Tween before being developed. The signal was developed using SuperSignal West Dura Extended Duration Substrate kit (ThermoFisher, Waltham, MA) and imaged using ChemiDoc Imaging (Bio-Rad, Hercules, CA). Western blot images were processed and analysed using Image Lab program (Bio-Rad, Hercules, CA). Quantification of band intensities were performed by Image Lab (Bio-Rad, Hercules, CA) with GAPDH or alpha-tubulin used to control protein loading. Samples were normalized to controls.

#### RNA extraction from whole zebrafish larvae and qPCR

To confirm *appa* and *appb* mRNA levels decrease in our double mutant (*appa^-/-^appb^-/-^*), RNA was extracted from 24 hpf whole larvae (10 larvae per n, n=5). Total RNA was extracted using TRI Reagent^®^ (Sigma, St. Louis, MO). Then, RNA samples were treated with RQ1 RNase-free DNase 1x reaction buffer and RQ1 RNase-free DNase (Promega, Fitchburg, WI). cDNA was synthesized using High-Capacity RNA-to-cDNA™ Kit (Applied Biosystems™, Waltham, MA) with RNase inhibitor and converted in a single-cycle reaction on a 2720 Thermal Cycler (Applied Biosystems™, Waltham, MA). Quantitative PCR was performed with inventoried TaqMan Gene Expression Assays with FAM reporter dye in TaqMan Universal PCR Master Mix with UNG (ThermoFisher, Waltham, MA). The assay was carried out on Micro-Amp 96- well optical microtiter plates (ThermoFisher, Waltham, MA) on a 7900HT Fast QPCR System (Applied Biosystems™, Waltham, MA). qPCR results were analysed with the SDS 2.3 software (Applied Biosystems™, Waltham, MA). cDNA values from each sample was normalized with average CT’s of house-keeping genes (*eef1a1l1* and *actb1*), then the relative quantity was determined using the ΔΔCT method (88) with the sample of wild-type sibling embryos (24 hpf) as the calibrator. TaqMan^®^ Gene Expression Assays (Applied Biosystems™, Waltham, MA) were used for the following genes: amyloid beta (A4) precursor protein A (*appa*) (Dr03144365_m1), eukaryotic translation elongation factor 1 alpha 1, like 1 (*eef1a1l1*) (Dr03432748_m1) and actin, beta 1 (*actb1*) (Dr03432610_m1).

#### Cilia length measurement in zebrafish larvae

To compare the number of brain ependymal cilia and their length, 30 hpf AB wild-type and *appa^-/-^appb^-/-^* zebrafish larvae were used. The larvae were treated with PTU, fixed in 4% PFA and the immunostaining with antibody against acetylated tubulin was performed as describe in the section above. Stacks (of around 25μm depending on the angle of the mounted sample) were taken in the region of interest (ROI) of the dorsal portion of the diencephalic ventricle using Zeiss LSM710 confocal microscope using inverted 40x water immersion objective (Plan-Apochromat 40x/1.0). Images were then processed using Imaris (BITPLANE^TM^, Belfast, United Kingdom) and the cilia length was measured with the acetylated tubulin signal using the “*measuring points*” tool of the program. Raw data of the measurement were exported to Microsoft Excel and compiled into GraphPad Prism® 7 for statistical analysis.

#### Brain ventricles injection and size measurement

To measure the size of the brain ventricles in live zebrafish, 2dpf PTU-treated zebrafish larvae were used. Rhodamine-Dextran injection protocol was performed as describe by Gutzman and Sive (89). Briefly, the larvae were anesthetized with tricaine in the EM and transferred onto a Petri dish covered with 1% agarose, lined with rows moulded. The larvae were kept in EM complemented with tricaine during the whole procedure and place on a ventral position, with top of their head facing upwards. Injections were performed using borosilicate injection needles previously pulled (P-97 Flaming/Brown micropipette puller) (Sutter Instrument, Novato, CA). Using a microinjector apparatus, 2nl of Rhodamine B isothiocyanate-Dextran (Sigma, St. Louis, MO) were injected in the hindbrain ventricle without perforating or hitting the brain tissue below.

Larvae with non-effective injections were sorted out using a fluorescent stereomicroscope (Nikon Instruments, Melville, NY). Quickly after the sorting, the larvae were mounted in 1% low-melting point agarose on glass bottom 35 mm Petri dish. Confocal imaging stacks were acquired using an inverted Nikon A1 confocal system using a 20x objective (Plan-Apochromat 20x/0,75). Image processing of the confocal stacks were done with Imaris program. The “*surface*” tool option was used for each sample. Data of the surface volume and area were automatically generated by the program. Length measurements of the areas of the ventricles were obtain manually with the “*measuring tool*”. All data were exported into Microsoft Excel and GraphPad® 7 Prism for statistical analysis.

#### Transmission electron microscopy

To evaluate the integrity of the internal structure of the axonemes and microtubules doublets of the brain motile cilia in older zebrafish, transmission electron microscopy was performed on fixed brains. Adult zebrafish were euthanized in tricaine and brains dissected, rinsed in ice-cold PBS and fixed in 2% PFA and 2% glutaraldehyde (Sigma, St. Louis, MO), in 0.042M Millonig buffer (0.081M Na_2_HPO_4_, 0.0183M NaH_2_PO_4_, 0.086M NaCl) pH 7.4 at least 24h at 4°C. After fixation, brains were cut in two halves and then treated in 2% osmium tetroxide (Sigma, St. Louis, MO) in 0.1M Millonig buffer pH 7.4. Specimens were then rinsed and incubated overnight in 4% sucrose solution in 0.1M Millonig buffer pH 7.4 after which they were dehydrated in series of ethanol and embedded in a mix of acetone and agar 100 resin plastic (TAAB Laboratories Equipment Ltd, Berks, United Kingdom) and allowed to polymerize for 48h. Blocks were trimmed as semi-thin (1 μm) and ultra-thin (70 nm) sections collected with a commercial ultramicrotome (Leica EM UC7, Leica Microsystems, Wetzlar, Germany). Sections were post-stained with 5% uranyl acetate in distilled H_2_O during 40-60 min, rinsed in distilled H_2_O and then treated with 0.3% Lead Citrate (ThermoFisher, Waltham, MA) for 30- 60 s. Images were acquired using secondary electron detection. Images were acquired with a Tecnai Spirit BT transmission electron microscope (Field Electron and Ion Company, Hillsboro, OR).

#### Statistical analysis

Statistical analysis was performed using GraphPad 7 software (Prism®, San Diego, CA). Data were presented as means with standard deviation (± SD) or standard errors of the mean (± SEM). For analysis of cilia length, D’Agostino & Pearson normality test (P < 0.0001) and non-parametric two-tailed Mann-Whitney U tests were performed. Results related to qPCR and western blot quantification, and ventricle size measurements were compared statistically using unpaired Student’s t-tests. Statistical significance was set at ρ < 0.05 (*), 0.01 (**), 0.005 (***) and 0.0001 (****).

## Acknowledgements

We thank Elisa Alexandersson and Katarina Türner Stenström for fish maintenance and the Centre for Cellular Imaging at the University of Gothenburg and the National Microscopy Infrastructure (VR-RFI 2016-00968) for microscopy support. We also thank Debora Kaminski for the mouse brains samples, Nathalie Jurish-Yaksi (Norwegian University of Science and Technology – NTNU) and Jean-François Papon (Public Hospital Network of Paris (AP-HP)) for insight and thoughtful discussions about cilia.

## Additional information

### Author contributions

**Jasmine Chebli:** Conceptualization, Formal analysis, Investigation, Visualization, Methodology, Data curation, Project administration, Writing - original draft, Writing - review and editing. **Maryam Rahmati**, **Tammaryn Lashley**, **Anders Oldfors** and **Birgitta Edeman**: Formal analysis, Writing - review and editing. **Henrik Zetterberg**: Resources, Supervision, Funding acquisition, Writing - original draft, Project administration, Writing - review and editing. **Alexandra Abramsson**: Conceptualization, Supervision, Formal analysis, Investigation, Visualization, Methodology, Writing - original draft, Project administration, Writing - review and editing. All authors reviewed and approved the final manuscript.

**Author ORCIDs:** JC: 0000-0003-0791-3198, TL: 0000-0001-7389-0348, AO: 0000-0002-5758-7397, HZ: 0000-0003-3930-4354, AA: 0000-0002-4715-9225.

### Ethics

All animal experiments in this study were performed in in accordance with the guidelines of the Swedish National Board for Laboratory Animals. Ethical approval for the use of human post-mortem samples was approved by a London Research Ethics Committee and tissue stored for research under a license from the Human Tissue Authority.

## Supplementary materials

**Supplementary file 1.**
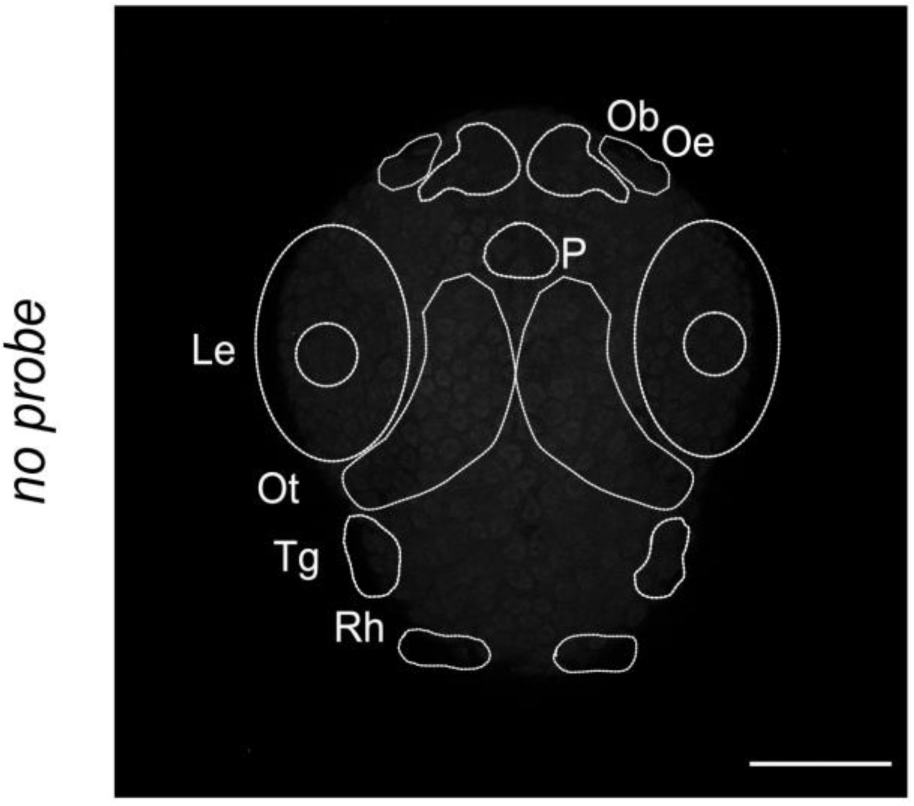
Negative control for whole-mount fluorescent *in situ*. Whole-mount fluorescent *in situ* in the absence of mRNA probe in 30 hpf WT larvae zebrafish. Maximum projection (77 stacks). T= telencephalic ventricle, D/M= diencephalic/mesencephalic ventricle, R= rhombencephalic ventricle, Ob= olfactory bulb, Oe= olfactory epithelium, P= pituitary gland, Le= lens, Ot= optic tectum, Tg= trigeminal ganglia, Rh= rhombomeres, Ov= otic vesicle. Magnification: 20x. Scale bar: 100µm.

**Supplementary file 2.**
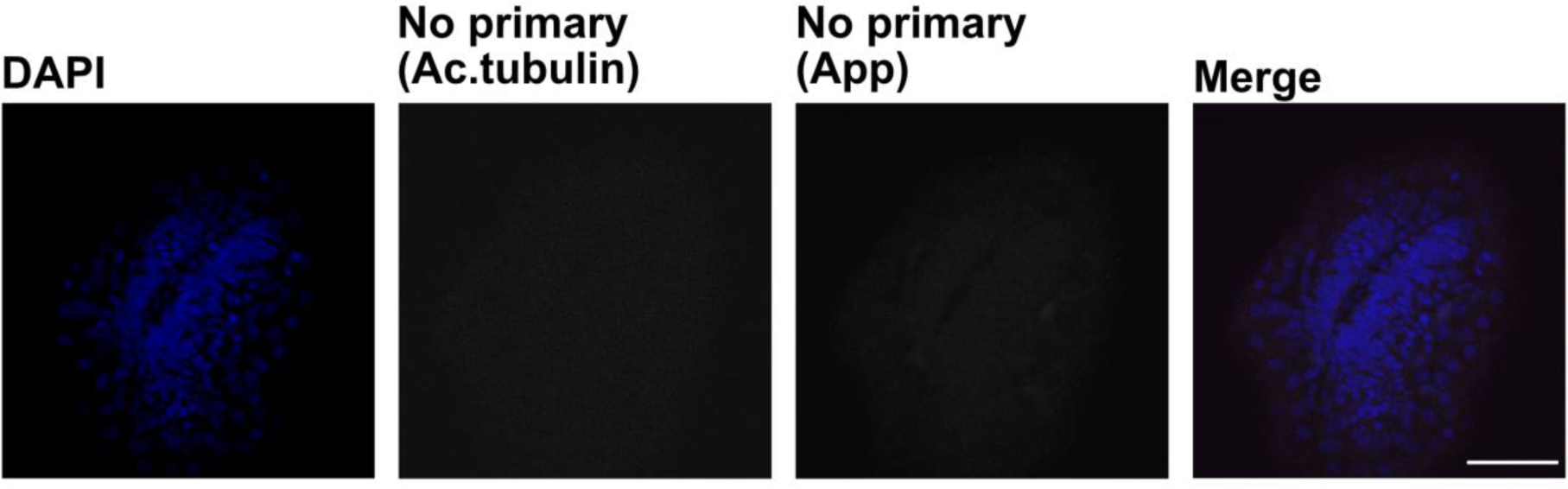
Negative controls for whole-mount fluroscent in situ. Negative controls for immunofluorescence in larvae zebrafish. Whole-mount immunofluorescence in larvae zebrafish with secondary antibodies without primary anti-acetylated tubulin and anti-App primary antibodies. Cell nuclei stained with DAPI (blue). Magnification: 40x. Scale bar: 50µm.

**Supplementary file 3.**
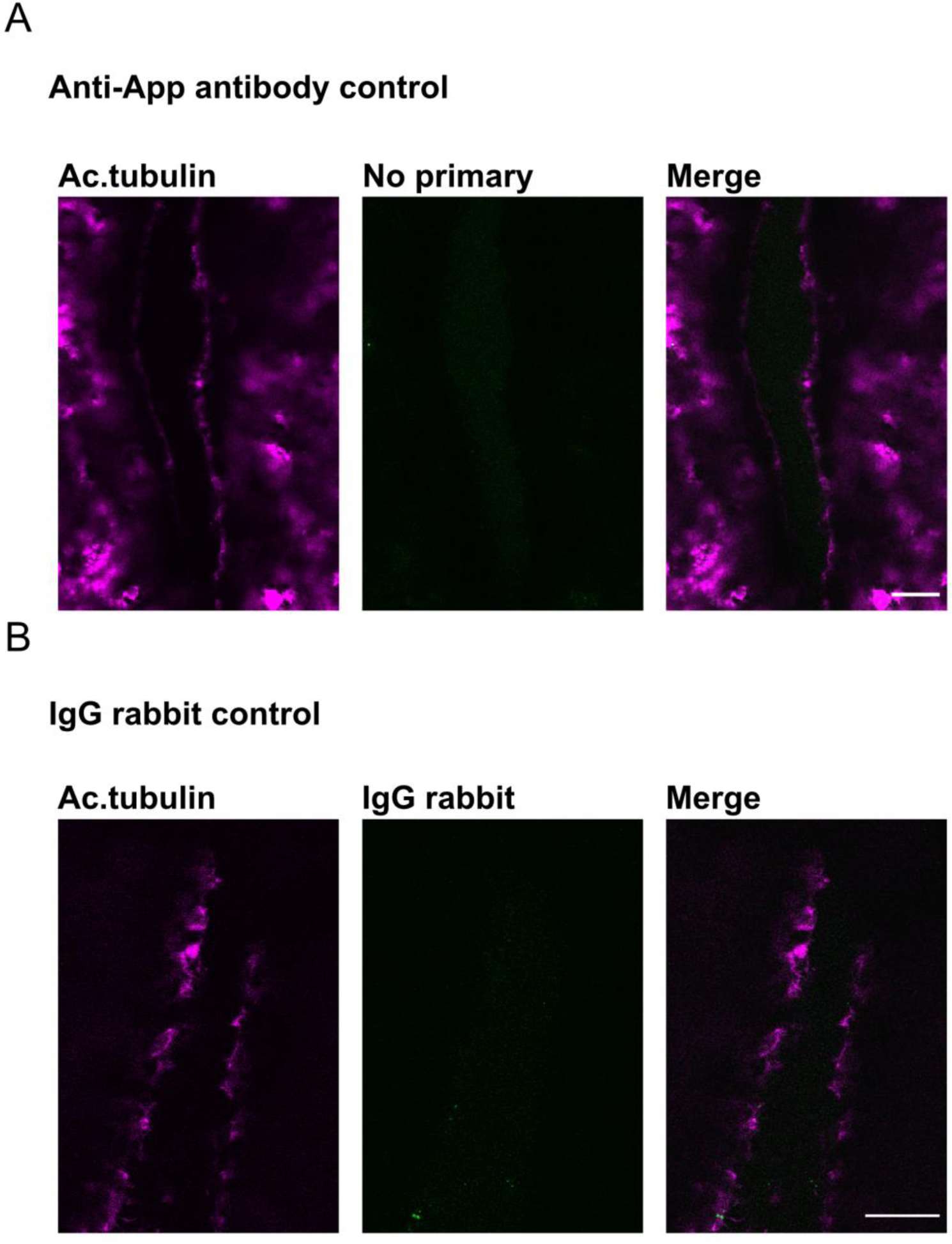
Negative controls for immunofluorescence in larvae zebrafish. Negative immunofluorescence control in adult zebrafish. Adult zebrafish brain slices stained with anti-acetylated tubulin antibody and (**A**) secondary anti-rabbit –Alexa488 antibody (without anti-App (Y188) antibody) or with (**B**) rabbit IgG serum and secondary anti-rabbit –Alexa488 antibody. Sec Magnification: (**A-B**)= 60x. Scale bar: (**A-B**)= 20µm.

**Supplementary file 4.**
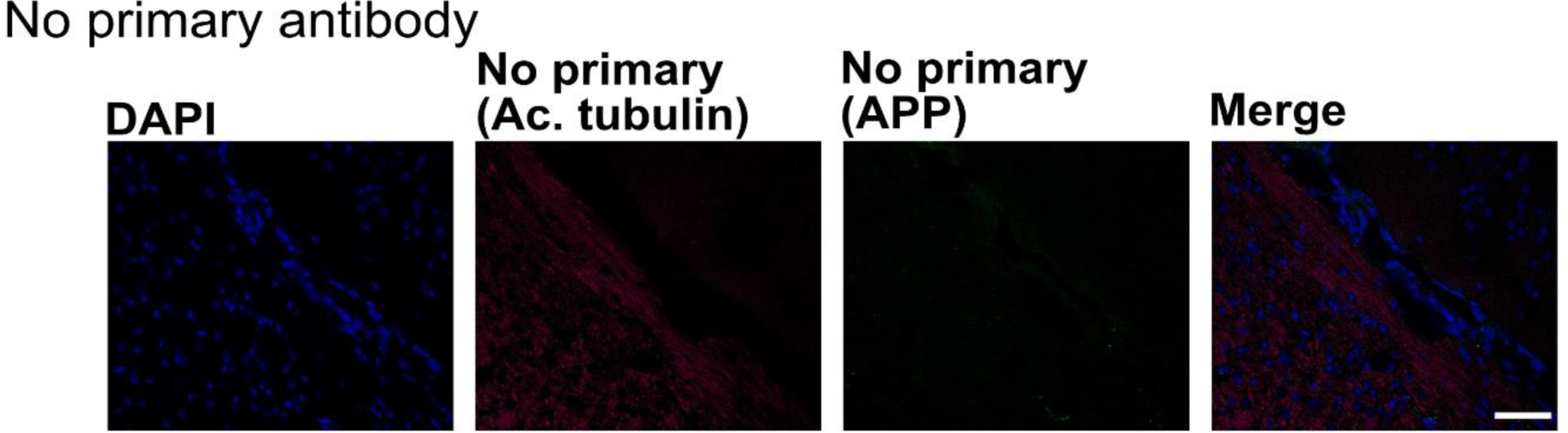
Negative immunostaining controls in adult mouse. Negative controls of Immunofluorescence staining on adult mouse brain. Slides incubated without primary antibodies and only with the corresponding secondary antibodies. For cell nuclei with DAPI (blue). Magnification: 40x. Scale bar: 50µm.

**Supplementary file 5.**
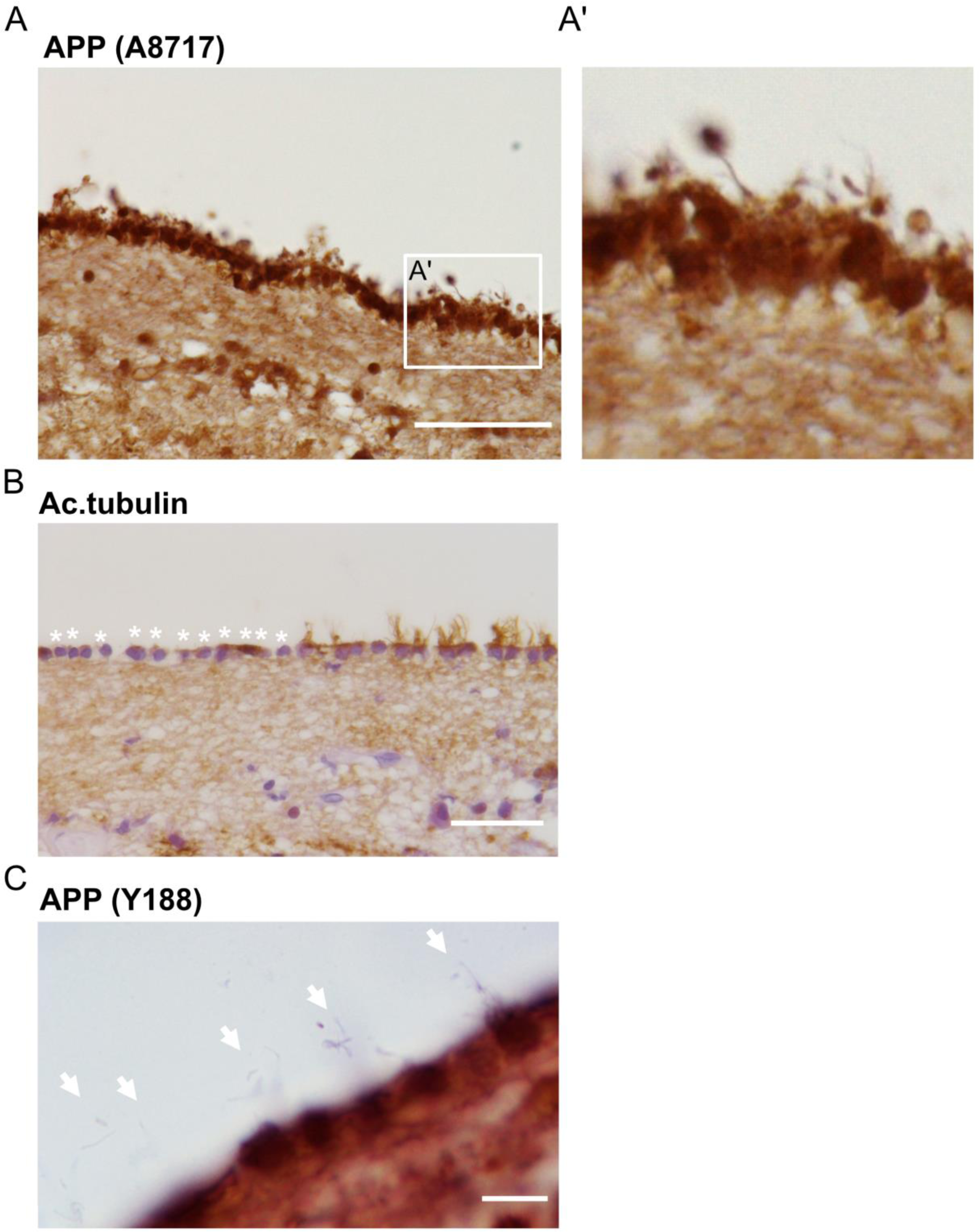
Immunostaining of human barin section with anti-APP (A8717) antibody and demaged cilia after tissue processing. Immunohistochemical staining of APP in human brain section. (**A**) Detection of APP with anti-APP 8A717 confirms the accumulation of APP within ependymal cells and along ependymal cilia. Close up in (**A’**). (**B**,**C**) Sections immunostained with an anti-acetylated tubulin (**B**) or an anti-APP (Y188) (**C**) antibody reveal that whereas some cilia seem to remain intact, some are completely damaged (see asterisks). White arrows indicate portions of cilia detached from their ependymal cells (**C**). Magnification: (**A,B**)= 40x, (**C**)= 100x. Scale bar: (**A**)= 50 µm, (**B**)= 10µm, (**C**)= 2µm.

**Supplementary file 7.**
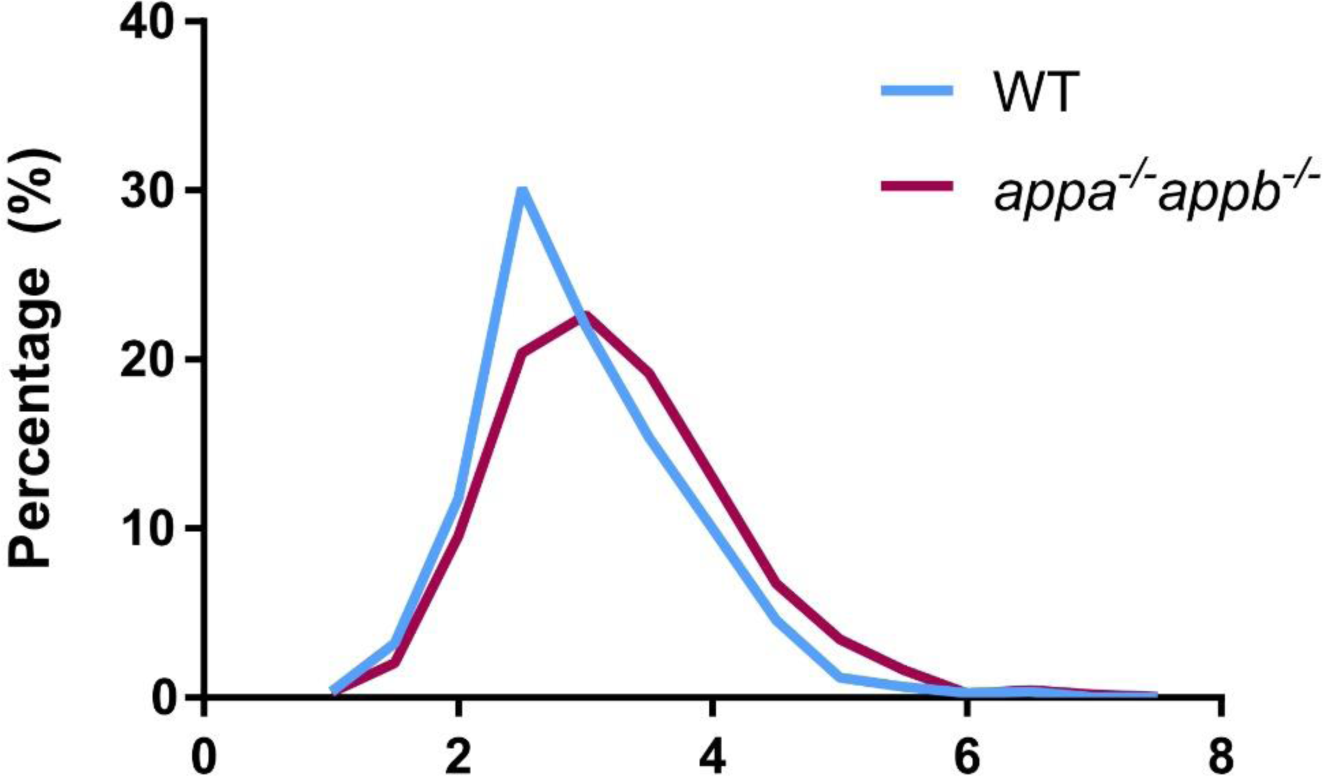
Frequency distribution of the length of brain ventricle in 30hpf larvae zebrafish. Frequency distribution of cilia length in 30hpf larvae zebrafish diencephalic/mesencephalic ventricle. A higher percentage of smaller cilia in WT (blue curve) compared to *appa^-/-^appb^-/-^* cilia population (magenta curve). n=10 WT (1091 cilia), n=16 *appa^-/-^appb^-/-^* (1511 cilia).

**Supplementary file 10.**
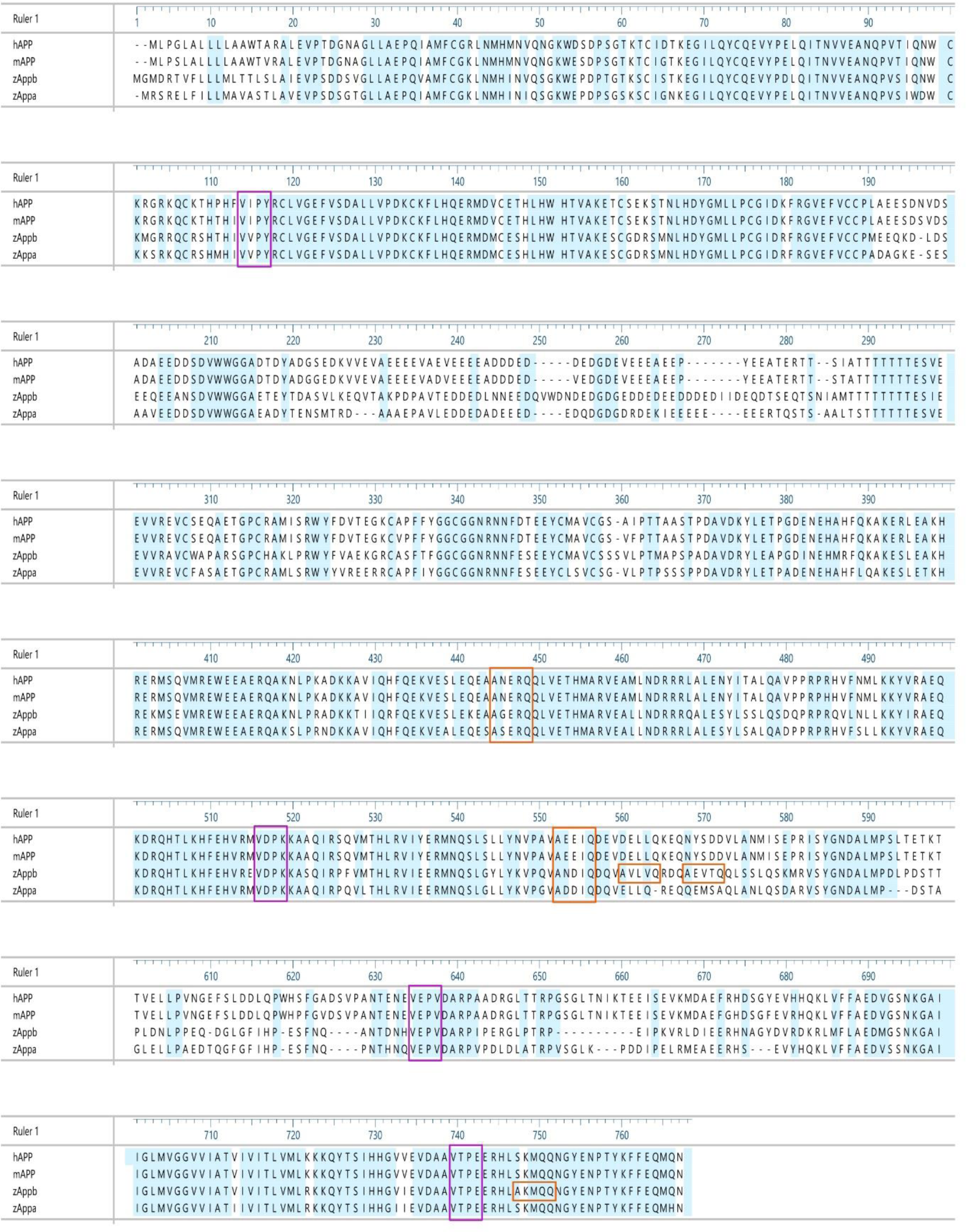
Cilia targeting sequences in human, mouse and zebrafish APP. Proteins sequence alignment of human APP751, mouse APP751 and zebrafish Appa738 and Appb751. Bright blue background shows conserved amino acids between species. Cilia targeting sequences AxxxQ (orange boxes) and VxPx (purple boxes).

## References

1. Selkoe DJ, Hardy J. The amyloid hypothesis of Alzheimer’s disease at 25 years. EMBO Mol Med. 2016;8(6):595–608.

2. Blennow K, de Leon MJ, Zetterberg H. Alzheimer’s disease. The Lancet. 2006;368(9533):387–403.

3. Wang B, Wang Z, Sun L, Yang L, Li H, Cole AL, et al. The Amyloid Precursor Protein Controls Adult Hippocampal Neurogenesis through GABAergic Interneurons. Journal of Neuroscience. 2014;34(40):13314–25.

4. Caille I, Allinquant B, Dupont E, Bouillot C, Langer A, Muller U, et al. Soluble form of amyloid precursor protein regulates proliferation of progenitors in the adult subventricular zone. Development. 2004;131(9):2173–81.

5. Young-Pearse TL, Chen AC, Chang R, Marquez C, Selkoe DJ. Secreted APP regulates the function of full-length APP in neurite outgrowth through interaction with integrin beta1. Neural Dev. 2008;3:15.

6. Deyts C, Thinakaran G, Parent AT. APP Receptor? To Be or Not To Be. Trends Pharmacol Sci. 2016;37(5):390–411.

7. Banote RK, Chebli J, Satir TM, Varshney GK, Camacho R, Ledin J, et al. Amyloid precursor protein-b facilitates cell adhesion during early development in zebrafish. Sci Rep. 2020;10(1):10127.

8. Wang Z, Wang B, Yang L, Guo Q, Aithmitti N, Songyang Z, et al. Presynaptic and postsynaptic interaction of the amyloid precursor protein promotes peripheral and central synaptogenesis. J Neurosci. 2009;29(35):10788–801.

9. Young-Pearse TL, Bai J, Chang R, Zheng JB, LoTurco JJ, Selkoe DJ. A Critical Function for -Amyloid Precursor Protein in Neuronal Migration Revealed by In Utero RNA Interference. Journal of Neuroscience. 2007;27(52):14459–69.

10. Muller UC, Deller T, Korte M. Not just amyloid: physiological functions of the amyloid precursor protein family. Nat Rev Neurosci. 2017;18(5):281–98.

11. Haass C, Kaether C, Thinakaran G, Sisodia S. Trafficking and proteolytic processing of APP. Cold Spring Harb Perspect Med. 2012;2(5):a006270.

12. Brown JM, Witman GB. Cilia and Diseases. Bioscience. 2014;64(12):1126–37.

13. Park SM, Jang HJ, Lee JH. Roles of Primary Cilia in the Developing Brain. Front Cell Neurosci. 2019;13:218.

14. Spassky N, Merkle FT, Flames N, Tramontin AD, Garcia-Verdugo JM, Alvarez- Buylla A. Adult ependymal cells are postmitotic and are derived from radial glial cells during embryogenesis. J Neurosci. 2005;25(1):10–8.

15. Lechtreck KF, Delmotte P, Robinson ML, Sanderson MJ, Witman GB. Mutations in Hydin impair ciliary motility in mice. J Cell Biol. 2008;180(3):633–43.

16. Abbott NJ, Pizzo ME, Preston JE, Janigro D, Thorne RG. The role of brain barriers in fluid movement in the CNS: is there a ’glymphatic’ system? Acta Neuropathol. 2018;135(3):387–407.

17. Ethell DW. Disruption of cerebrospinal fluid flow through the olfactory system may contribute to Alzheimer’s disease pathogenesis. J Alzheimers Dis. 2014;41(4):1021–30.

18. Kapoor KG, Katz SE, Grzybowski DM, Lubow M. Cerebrospinal fluid outflow: an evolving perspective. Brain Res Bull. 2008;77(6):327–34.

19. Klein S, Goldman A, Lee H, Ghahremani S, Bhakta V, Center UCG, et al. Truncating mutations in APP cause a distinct neurological phenotype. Ann Neurol. 2016;80(3):456–60.

20. Kiprilov EN, Awan A, Desprat R, Velho M, Clement CA, Byskov AG, et al. Human embryonic stem cells in culture possess primary cilia with hedgehog signaling machinery. J Cell Biol. 2008;180(5):897–904.

21. Magara F, Muller U, Li ZW, Lipp HP, Weissmann C, Stagljar M, et al. Genetic background changes the pattern of forebrain commissure defects in transgenic mice underexpressing the -amyloid-precursor protein. Proceedings of the National Academy of Sciences. 1999;96(8):4656–61.

22. Baudoin JP, Viou L, Launay PS, Luccardini C, Espeso Gil S, Kiyasova V, et al. Tangentially migrating neurons assemble a primary cilium that promotes their reorientation to the cortical plate. Neuron. 2012;76(6):1108–22.

23. Guo J, Otis JM, Higginbotham H, Monckton C, Cheng J, Asokan A, et al. Primary Cilia Signaling Shapes the Development of Interneuronal Connectivity. Dev Cell. 2017;42(3):286–300 e4.

24. Higginbotham H, Guo J, Yokota Y, Umberger NL, Su CY, Li J, et al. Arl13b- regulated cilia activities are essential for polarized radial glial scaffold formation. Nat Neurosci. 2013;16(8):1000–7.

25. Guo J, Otis JM, Suciu SK, Catalano C, Xing L, Constable S, et al. Primary Cilia Signaling Promotes Axonal Tract Development and Is Disrupted in Joubert Syndrome-Related Disorders Models. Developmental Cell. 2019;51(6):759–74.e5.

26. Galati DF, Sullivan KD, Pham AT, Espinosa JM, Pearson CG. Trisomy 21 Represses Cilia Formation and Function. Dev Cell. 2018;46(5):641–50 e6.

27. Chakravarthy B, Gaudet C, Ménard M, Brown L, Atkinson T, LaFerla FM, et al. Reduction of the immunostainable length of the hippocampal dentate granule cells’ primary cilia in 3xAD-transgenic mice producing human Aβ1-42 and tau. Biochemical and Biophysical Research Communications. 2012;427(1):218–22.

28. Ibanez-Tallon I, Pagenstecher A, Fliegauf M, Olbrich H, Kispert A, Ketelsen UP, et al. Dysfunction of axonemal dynein heavy chain Mdnah5 inhibits ependymal flow and reveals a novel mechanism for hydrocephalus formation. Hum Mol Genet. 2004;13(18):2133–41.

29. Vorobyeva AG, Saunders AJ. Amyloid-beta interrupts canonical Sonic hedgehog signaling by distorting primary cilia structure. Cilia. 2018;7:5.

30. Musa A, Lehrach H, Russo VA. Distinct expression patterns of two zebrafish homologues of the human APP gene during embryonic development. Dev Genes Evol. 2001;211(11):563–7.

31. Tanimoto M, Ota Y, Inoue M, Oda Y. Origin of inner ear hair cells: morphological and functional differentiation from ciliary cells into hair cells in zebrafish inner ear. J Neurosci. 2011;31(10):3784–94.

32. Kawarabayashi T, Shoji M, Harigaya Y, Yamaguchi H, Hirai S. Amyloid beta/A4 protein precursor is widely distributed in both the central and peripheral nervous systems of the mouse. Brain Res. 1991;552(1):1–7.

33. Ohta M, Kitamoto T, Iwaki T, Ohgami T, Fukui M, Tateishi J. Immunohistochemical distribution of amyloid precursor protein during normal rat development. Brain Res Dev Brain Res. 1993;75(2):151–61.

34. Stern RA, Otvos L, Jr., Trojanowski JQ, Lee VM. Monoclonal antibodies to a synthetic peptide homologous with the first 28 amino acids of Alzheimer’s disease beta-protein recognize amyloid and diverse glial and neuronal cell types in the central nervous system. Am J Pathol. 1989;134(5):973–8.

35. Fame RM, Chang JT, Hong A, Aponte-Santiago NA, Sive H. Directional cerebrospinal fluid movement between brain ventricles in larval zebrafish. Fluids Barriers CNS. 2016;13(1):11.

36. Manton I.; Clarke B. An electron microscope study of the spermatozoid of sphagnum. Journal of Experimental Botany. 1952;3:265–75.

37. Satir P. Studies on cilia: II. Examination of the distal region of the ciliary shaft and the role of the filaments in motility. Journal of Cell Biology. 1965(26):805–34.

38. Satir P, Heuser T, Sale WS. A Structural Basis for How Motile Cilia Beat. Bioscience. 2014;64(12):1073–83.

39. Tam BM, Moritz OL, Hurd LB, Papermaster DS. Identification of an outer segment targeting signal in the COOH terminus of rhodopsin using transgenic Xenopus laevis. J Cell Biol. 2000;151(7):1369–80.

40. Deretic D. A role for rhodopsin in a signal transduction cascade that regulates membrane trafficking and photoreceptor polarity. Vision Research. 2006;46(27):4427–33.

41. Rakoczy EP, Kiel C, McKeone R, Stricher F, Serrano L. Analysis of disease- linked rhodopsin mutations based on structure, function, and protein stability calculations. J Mol Biol. 2011;405(2):584–606.

42. Berbari NF, Johnson AD, Lewis JS, Askwith CC, Mykytyn K. Identification of ciliary localization sequences within the third intracellular loop of G protein-coupled receptors. Mol Biol Cell. 2008;19(4):1540–7.

43. Domire JS, Green JA, Lee KG, Johnson AD, Askwith CC, Mykytyn K. Dopamine receptor 1 localizes to neuronal cilia in a dynamic process that requires the Bardet-Biedl syndrome proteins. Cell Mol Life Sci. 2011;68(17):2951–60.

44. Chakravarthy B, Gaudet C, Menard M, Atkinson T, Chiarini A, Dal Pra I, et al. The p75 neurotrophin receptor is localized to primary cilia in adult murine hippocampal dentate gyrus granule cells. Biochem Biophys Res Commun. 2010;401(3):458–62.

45. Ye F, Breslow DK, Koslover EF, Spakowitz AJ, Nelson WJ, Nachury MV. Single molecule imaging reveals a major role for diffusion in the exploration of ciliary space by signaling receptors. eLife. 2013;2.

46. de Coninck D, Schmidt TH, Schloetel JG, Lang T. Packing Density of the Amyloid Precursor Protein in the Cell Membrane. Biophys J. 2018;114(5):1128–41.

47. Yang J, Li T. The ciliary rootlet interacts with kinesin light chains and may provide a scaffold for kinesin-1 vesicular cargos. Exp Cell Res. 2005;309(2):379–89.

48. Long H, Huang K. Transport of Ciliary Membrane Proteins. Frontiers in Cell and Developmental Biology. 2020;7.

49. Malicki J, Avidor-Reiss T. From the cytoplasm into the cilium: bon voyage. Organogenesis. 2014;10(1):138–57.

50. Mita S, Schon EA, Herbert J. Widespread expression of amyloid beta-protein precursor gene in rat brain. Am J Pathol. 1989;134(6):1253–61.

51. Alvarez-Buylla A, Lim DA. For the long run: maintaining germinal niches in the adult brain. Neuron. 2004;41(5):683–6.

52. Zhang X, Jia S, Chen Z, Chong YL, Xie H, Feng D, et al. Cilia-driven cerebrospinal fluid flow directs expression of urotensin neuropeptides to straighten the vertebrate body axis. Nat Genet. 2018;50(12):1666–73.

53. Song Z, Zhang X, Jia S, Yelick PC, Zhao C. Zebrafish as a Model for Human Ciliopathies. J Genet Genomics. 2016;43(3):107–20.

54. Lowery LA, De Rienzo G, Gutzman JH, Sive H. Characterization and classification of zebrafish brain morphology mutants. Anat Rec (Hoboken). 2009;292(1):94–106.

55. Lowery LA, Sive H. Initial formation of zebrafish brain ventricles occurs independently of circulation and requires the nagie oko and snakehead/atp1a1a.1 gene products. Development. 2005;132(9):2057–67.

56. Olstad EW, Ringers C, Hansen JN, Wens A, Brandt C, Wachten D, et al. Ciliary Beating Compartmentalizes Cerebrospinal Fluid Flow in the Brain and Regulates Ventricular Development. Curr Biol. 2019;29(2):229–41 e6.

57. Morimoto Y, Yoshida S, Kinoshita A, Satoh C, Mishima H, Yamaguchi N, et al. Nonsense mutation in CFAP43 causes normal-pressure hydrocephalus with ciliary abnormalities. Neurology. 2019;92(20):e2364–e74.

58. Sawamoto K. New Neurons Follow the Flow of Cerebrospinal Fluid in the Adult Brain. Science. 2006;311(5761):629–32.

59. Demars M, Hu YS, Gadadhar A, Lazarov O. Impaired neurogenesis is an early event in the etiology of familial Alzheimer’s disease in transgenic mice. J Neurosci Res. 2010;88(10):2103–17.

60. Ma QH, Bagnard D, Xiao ZC, Dawe GS. A TAG on to the neurogenic functions of APP. Cell Adh Migr. 2008;2(1):2–8.

61. Giacomini A, Stagni F, Trazzi S, Guidi S, Emili M, Brigham E, et al. Inhibition of APP gamma-secretase restores Sonic Hedgehog signaling and neurogenesis in the Ts65Dn mouse model of Down syndrome. Neurobiol Dis. 2015;82:385–96.

62. Clement A, Solnica-Krezel L, Gould KL. The Cdc14B phosphatase contributes to ciliogenesis in zebrafish. Development. 2011;138(2):291–302.

63. Fame RM, Lehtinen MK. Emergence and Developmental Roles of the Cerebrospinal Fluid System. Dev Cell. 2020;52(3):261–75.

64. Zappaterra MW, Lehtinen MK. The cerebrospinal fluid: regulator of neurogenesis, behavior, and beyond. Cell Mol Life Sci. 2012;69(17):2863–78.

65. LeMay M, Alvarez N. The relationship between enlargement of the temporal horns of the lateral ventricles and dementia in aging patients with Down syndrome. Neuroradiology. 1990;32(2):104–7.

66. Ezratty EJ, Pasolli HA, Fuchs E. A Presenilin-2-ARF4 trafficking axis modulates Notch signaling during epidermal differentiation. J Cell Biol. 2016;214(1):89–101.

67. Nager AR, Goldstein JS, Herranz-Perez V, Portran D, Ye F, Garcia-Verdugo JM, et al. An Actin Network Dispatches Ciliary GPCRs into Extracellular Vesicles to Modulate Signaling. Cell. 2017;168(1-2):252–63 e14.

68. Perez-Gonzalez R, Gauthier SA, Kumar A, Levy E. The exosome secretory pathway transports amyloid precursor protein carboxyl-terminal fragments from the cell into the brain extracellular space. J Biol Chem. 2012;287(51):43108–15.

69. Spitzer P, Mulzer LM, Oberstein TJ, Munoz LE, Lewczuk P, Kornhuber J, et al. Microvesicles from cerebrospinal fluid of patients with Alzheimer’s disease display reduced concentrations of tau and APP protein. Sci Rep. 2019;9(1):7089.

70. Leinonen V, Kuulasmaa T, Hiltunen M. iNPH-the mystery resolving. EMBO Mol Med. 2021;13(3):e13720.

71. Jeppsson A, Zetterberg H, Blennow K, Wikkelsø C. Idiopathic normal-pressure hydrocephalus: pathophysiology and diagnosis by CSF biomarkers. Neurology. 2013;80(15):1385–92.

72. Jeppsson A, Holtta M, Zetterberg H, Blennow K, Wikkelso C, Tullberg M. Amyloid mis-metabolism in idiopathic normal pressure hydrocephalus. Fluids Barriers CNS. 2016;13(1):13.

73. Pyykko OT, Lumela M, Rummukainen J, Nerg O, Seppala TT, Herukka SK, et al. Cerebrospinal fluid biomarker and brain biopsy findings in idiopathic normal pressure hydrocephalus. PLoS One. 2014;9(3):e91974.

74. Kim JY, Rasheed A, Yoo SJ, Kim SY, Cho B, Son G, et al. Distinct amyloid precursor protein processing machineries of the olfactory system. Biochem Biophys Res Commun. 2018;495(1):533–8.

75. Reiten I, Uslu FE, Fore S, Pelgrims R, Ringers C, Diaz Verdugo C, et al. Motile-Cilia-Mediated Flow Improves Sensitivity and Temporal Resolution of Olfactory Computations. Curr Biol. 2017;27(2):166–74.

76. Yoshihara Y. Zebrafish Olfactory System. Springer Japan; 2014. p. 71–96.

77. Doty RL. The olfactory system and its disorders. Semin Neurol. 2009;29(1):74–81.

78. Trudeau S, Anne S, Otteson T, Hopkins B, Georgopoulos R, Wentland C. Diagnosis and patterns of hearing loss in children with severe developmental delay. Am J Otolaryngol. 2021;42(3):102923.

79. Liu Y, Fang S, Liu LM, Zhu Y, Li CR, Chen K, et al. Hearing loss is an early biomarker in APP/PS1 Alzheimer’s disease mice. Neurosci Lett. 2020;717:134705.

80. Omata Y, Tharasegaran S, Lim Y-M, Yamasaki Y, Ishigaki Y, Tatsuno T, et al. Expression of amyloid-β in mouse cochlear hair cells causes an early-onset auditory defect in high-frequency sound perception. Aging. 2016;8(3):427–39.

81. Westerfield M. The Zebrafish Book : A Guide for the Laboratory Use of Zebrafish. http://zfinorg/zf_info/zfbook/zfbkhtml. 2000.

82. Varshney GK, Carrington B, Pei W, Bishop K, Chen Z, Fan C, et al. A high-throughput functional genomics workflow based on CRISPR/Cas9-mediated targeted mutagenesis in zebrafish. Nature protocols. 2016;11(12):2357–75.

83. UniProt C. UniProt: the universal protein knowledgebase in 2021. Nucleic Acids Res. 2021;49(D1):D480–D9.

84. Banote RK, Chebli J, Şatır TM, Varshney GK, Camacho R, Ledin J, et al. Amyloid precursor protein-b facilitates cell adhesion during early development in zebrafish. Sci Rep. 2020;10(1):10127.

85. Kimmel CB, Ballard WW, Kimmel SR, Ullmann B, Schilling TF. Stages of embryonic development of the zebrafish. Dev Dyn. 1995;203(3):253–310.

86. Lauter G, Söll I, Hauptmann G. Sensitive whole-mount fluorescent in situ hybridization in zebrafish using enhanced tyramide signal amplification. Methods Mol Biol. 2014;1082:175–85.

87. Lashley T, Rohrer JD, Bandopadhyay R, Fry C, Ahmed Z, Isaacs AM, et al. A comparative clinical, pathological, biochemical and genetic study of fused in sarcoma proteinopathies. Brain. 2011;134:2548–64.

88. Livak KJ, Schmittgen TD. Analysis of relative gene expression data using real-time quantitative PCR and the 2(-Delta Delta C(T)) Method. Methods (San Diego, Calif). 2001;25(4):402–8.

89. Gutzman JH, Sive H. Zebrafish brain ventricle injection. J Vis Exp. 2009(26).

